# Dual Role of Cx43 in Enhancing CDK4/6 Inhibitor Response and Promoting the Elimination of Therapy-Induced Senescent Tumor Cells

**DOI:** 10.64898/2026.10.26.761820

**Authors:** Paula Carpintero-Fernández, Marta Barturen-Gómez, Vanesa Álvarez, Alexander Carneiro-Figueira, Soumadipta Mazumdar, Loretta De Chiara, Alejandro García-Yuste, MF Soler, Susana B. Bravo-López, Joaquín Mosquera, Benigno Acea, Carlos Fernandez-Lozano, María M. Caffarel, Constance Alabert, María D. Mayán

**Author notes:** Corresponding author: María D. Mayán. CellCOM Research Group. Center for Research in Nanomaterials and Biomedicine (CINBIO) and Institute of Biomedical Research of Ourense-Pontevedra-Vigo (IBI), University of Vigo. The Galician Health Service, SERGAS. Olimpia Valencia building, Campus Universitario Lagoas Marcosende, 36310, Pontevedra, Spain.

## Abstract

Breast cancer has become the most prevalent malignancy worldwide. A key clinical advance has been the introduction of cyclin-dependent kinase 4/6 inhibitors (CDK4/6i), that in combination with endocrine therapy has been a major advance for the treatment of luminal (ER⁺/HER2⁻) breast cancer. CDK4/6i induce cell cycle arrest in G1 and cellular senescence improving clinical outputs. However, limited efficacy and therapeutic resistance remain major challenges due to incomplete effects or the failure to eliminate the accumulated tumor senescent cells. Here, we identify connexin 43 (Cx43), a membrane channel protein involved in cell-to-cell communication, identified as a novel target in luminal breast cancer that potentiates the effects of CDK4/6 inhibitors. We found that Cx43 not only reduces tumor cell proliferation by enhancing senescence but also reshapes the Senescence-Associated Secretory Phenotype (SASP) profile. We also demonstrate that, in combination with CDK4/6i, Cx43 reinforces G1 arrest via transcriptional downregulation of E2F targets including CDK4 and CDK6, and through interaction with p21/p27, deepens the senescent state. Importantly, the Cx43-associated senescent phenotype sensitizes tumor cells accumulated under CDK4/6i treatment to the senolytic agent Navitoclax. Our results demonstrate that upregulation of Cx43 in breast cancer reduces proliferation and enhances the clearance of therapy-induced senescent cells. The triple-drug strategy comprising CDK4/6i, Cx43 delivery, and a senolytic drug such as Navitoclax holds great promise for overcoming CDK4/6i resistance and improving clinical outcomes in ER⁺/HER2⁻ breast cancer and may also be applicable to other cancer types that respond to CDK4/6i or other pro-senescence antitumoral drugs.

## INTRODUCTION

Breast cancer (BC) is the most diagnosed cancer worldwide and represents the fifth cause of cancer mortality globally [1] [2]. Although treatment of primary breast cancer has significantly improved in recent years, the prognosis for metastatic or recurrent disease remains poor. For instance, patients with estrogen receptor-positive (ER+)/human epidermal growth factor receptor 2-negative (HER2-) metastatic breast cancer have a median overall survival ranging from approximately 42 to 64 months, depending on the specific treatment regimen, with a five-year relative survival rate for this subtype around 31% [3]. Despite advancements in treatment options, the overall outlook for these patients remains challenging [1, 2]. In the last few years, CDK4/6i (Palbociclib, Abemaciclib, Ribociclib), a class of new-generation therapeutics, have become a revolution in the field of cancer and, in particular, in metastatic breast cancer; improving progression-free survival (PFS) and the overall survival (OS) in around 50 months in the case of ER+/HER2-breast cancer patients [4, 5] which is the most diagnosed breast cancer subtype. The three inhibitors are standard of care in luminal metastatic ER+/HER2-breast cancer combined with hormonotherapy, both in first- and second-line therapy with high response rates compared to hormone therapy alone [4, 6, 7]. They are approved to treat breast cancer and are also undergoing clinical trials against other tumor types [8].Their main mechanism of action is based on the induction of a stable cell cycle arrest through the inhibition of the Rb-CDK4/6 signalling pathway and the induction of a senescent-like phenotype in cancer cells [9–11]. However, and despite significant efforts in recent years to elucidate the full mechanisms of action of CDK4/6i, the specific pathways and factors influencing their response, efficacy, and resistance remain a substantial knowledge gap, limiting their therapeutic potential [10, 12–16]. Notably, subpopulations of tumor cells that enter a senescent-like state have been reported, under certain conditions, to re-enter the cell cycle, acquire stem cell-like properties, and contribute to disease recurrence [17]. To address this challenge, a therapeutic strategy known as the "one-two punch" has been proposed [18, 19] where senescence is first induced by pro-senescent agents such as CDK4/6i and subsequently targeted by senolytic drugs to eliminate senescent tumor cells. This approach has shown promising results [12,13], however CDK4/6i-treated breast cancer cells were reported to exhibit various senescence-associated phenotypes, and remain insensitive to common senolytic compounds [20]. Recent studies are beginning to uncover distinct mechanisms governing the senolytic response of these therapy-induced senescent tumor cells, highlighting the complexity of their clearance[20].

Connexins (Cxs) are channel-forming proteins involved in cell-to-cell communication that modulate cell growth and the tumor microenvironment via hemichannels, gap junctions (GJ), extracellular vesicles (EVs), tunnelling nanotubes or via channel-independent functions regulating different cellular signalling cascades [21–23]. Different reports suggest that Cxs act as tumor suppressors as their re-expression in tumor cells decrease their tumorigenicity and reverses the transformed phenotypes of the tumor cells [24] [25] [23, 26] particularly Cx43 [27] [28] [29, 30]. However, their role depends on the type of connexin, the type of tumor, and the specific microenvironmental context [31]. Connexin43 (Cx43) has been proposed as a favourable prognostic marker in breast cancer [32], with expression shown to decline progressively with increasing malignancy [33], however no studies have deepened its role in ER⁺/HER2⁻. Moreover, connexins and gap junctions have been associated with immune-mediated anti-tumor responses [34] and the regulation of cell cycle [35] [36] [37] [24] [27]. Notably, we have previously shown that Cx43 promotes cellular senescence in both non-tumoral [38] and tumoral contexts [30]. Also, recent findings from our group demonstrate that upregulation of Cx43 in BRAF-mutant tumors not only enhances senescence but also disrupts DNA repair mechanisms, increasing genomic instability and thereby contributing to improved therapeutic efficacy[30].

In this study we have investigated the role of Cx43 in ER+/HER2-human BC models and in response to CDK4/6i treatment. Our results demonstrated that the restoration of Cx43 in ER+/HER2-BC cells induce a reduction in proliferation and cell cycle arrest in G1 phase together with an increase in cellular senescence. The combination of Cx43 with CDK4/6i significantly increases the efficacy of the inhibitors. In this context, Cx43 contributes to increased cell death in the presence of the senolytic drug Navitoclax. Based on these results, we developed a system for delivering Cx43 mRNA and protein to tumor cells, successfully restoring Cx43 activity in ER⁺/HER2⁻ breast cancer cells. Altogether, our results identify a novel target with antitumor activity in ER⁺/HER2⁻ breast cancer cells that enhances CDK4/6 inhibitor efficacy and demonstrate a new and effective combination strategy based on restoring Cx43 in breast cancer cells using extracellular vesicles (EVs) as delivery vectors.

## RESULTS

### Cx43 Reduces Proliferation and Promotes a Senescent-Like Phenotype with a Specific SASP Signature in ER+/HER2-Breast Cancer Cells

To study the role of Cx43 in ER+/HER2-BC subtype, we took advantage of the human MCF7 cell line to analyze the expression and protein levels of different connexins and pannexins including Cx43 (Supplementary Figure S1a). Cx43 was hardly detected in MCF7 ER+/HER2-BC cells by RT-PCR and immunofluorescence (IF) (Figure 1a). This finding was confirmed by immunohistochemistry in ER⁺/HER2⁻ tumor and lymph node samples from patients, where Cx43 was undetectable or showed only weak, diffuse cytoplasmic staining (Figure 1b). These results are in line with those obtained from METABRIC database, where Cx43 mRNA is found to be downregulated in the tumoral area and low levels of Cx43 (mRNA) are associated with a recurrence in the disease (Figure S1b). Patient overall survival (OS) probability also correlates with higher levels of Cx43 mRNA (Figure 1c), considering, based on these results, Cx43 as a sign of good prognosis in ER+/HER2-BC subtype.

**Figure 1.**
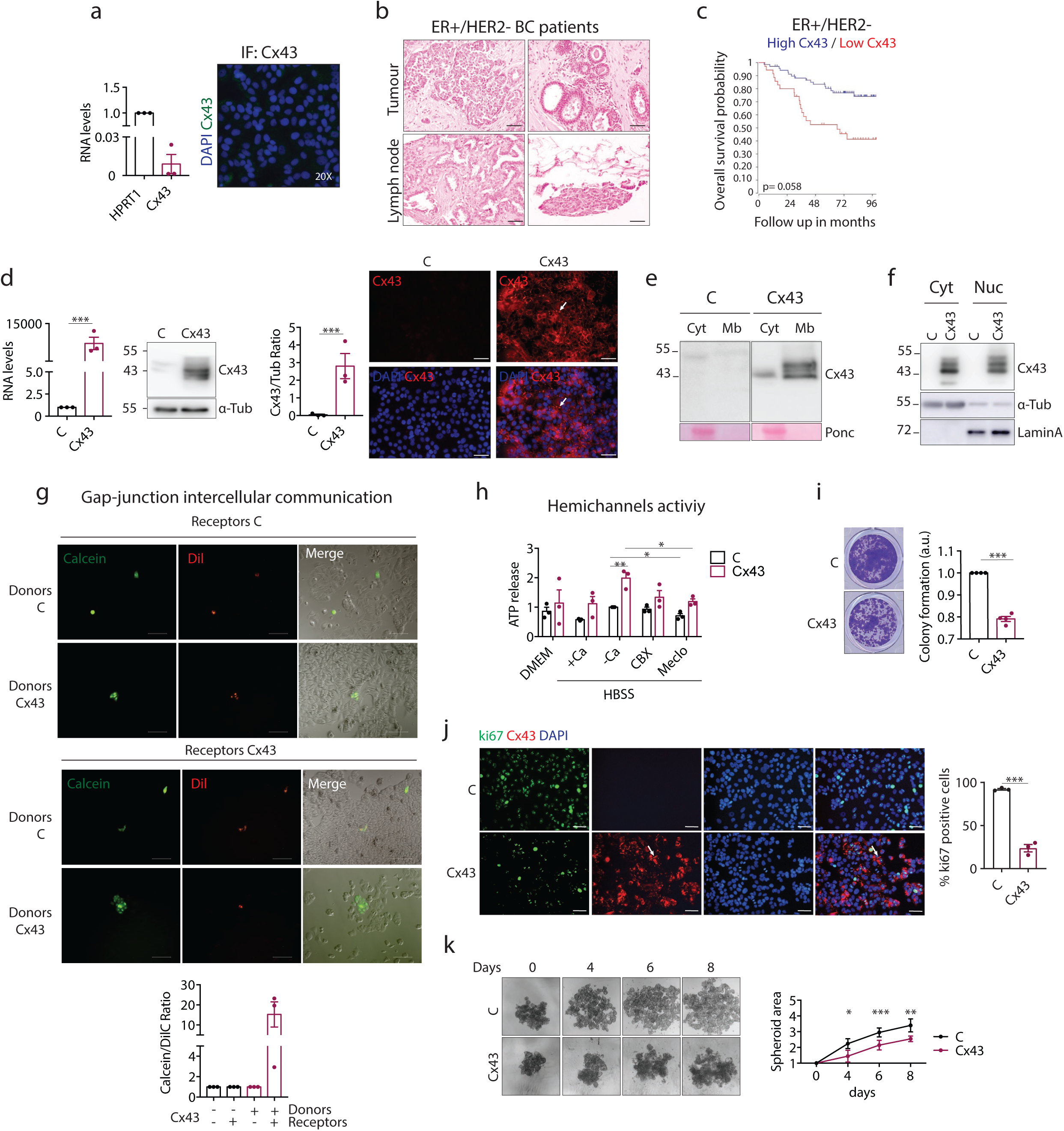
Cx43 induces proliferation arrest and a senescent-like phenotype in ER+/HER2- breast cancer cells. (a) Cx43 protein and mRNA levels in MCF7 ER+/HER2-breast cancer cells. (b) Cx43 immunohistochemistry from ER+/HER2-breast cancer patients from tumoral and lymph node areas. Images show 2 representative patients. Scale bar: 50 µM (c) Overall survival probability of breast cancer patients with high- and low-level expression of Cx43. The data have been obtained using the free online available tool Kapplan-Meier plotter. (d) ER+/HER2-breast cancer cells (MCF7) were infected with a plasmid to ectopically express Cx43. Cx43 protein and mRNA levels were tested by Western-blot (protein) and qPCR (mRNA) comparing the vector control (C) to MCF7-Cx43 cells. Tubulin was used as loading control. Immunofluorescence showing Cx43 expression and localization in MCF7 cells. White arrow indicating Cx43 gap junction plaques at the plasma membrane. Scale bar: 50 µM. (e) Western-blot demonstrating that Cx43 is localized at the plasma membrane in MCF7 cells after Cx43 expression compared to the C. Ponceau was used as loading control. (f) Western-blot demonstrating that Cx43 is located both in the cytoplasm and the nuclei in MCF7 cells after Cx43 expression compared to the C. Tubulin and LaminA were used as loading controls. (g) Parachute assay to confirm gap-junction intercellular communication. Scale bar: 50 µM. Quantification of parachute assay is shown in the panel below. (h) ATP release assay showing that Cx43 expression increased the hemichannel activity of MCF7 cells. Carbenoxolone and Meclofenamate were used as hemichannel inhibitors. (i) Colony formation assay showing a reduction in proliferation after Cx43 expression in ER+/HER2-BC cells. (j) Co-Immunofluorescence showing the correlation between Cx43 (red) expression and the cell proliferation marker Ki67 (green) in MCF7 C and Cx43-positive cells. White arrow indicating Cx43 gap junction plaques at the plasma membrane. Scale bar: 50 µM. (k) Spheroids of MCF7 cells (C and Cx43) to demonstrate cell proliferation arrest in a 3D culture. Spheroids were cultured for 8 days. The graph represents the mean of 6 spheroids/day. Data represent the mean ± SEM or SD (in the case of spheroids area) of 3 independent experiments. Two-tailed Student’s t-test and ANOVA were used to calculate the significance represented as followed: *p<0.05; **p<0.01; ***p<0.001.

Next, we decided to restore Cx43 in human ER+/ER2-BC cells by using a retroviral vector (Figure 1d). Since connexins activity and function depend on their subcellular localization, we checked Cx43 localization by using immunofluorescence (Figure 1d) and western blot (Figure 1e, f). Figure 1d shows that in Cx43-restored cells, Cx43 (in red) predominantly localized at cell– cell contacts (white arrow). Western blot analysis of cytoplasmic (soluble) and membrane (insoluble) cellular fractions confirmed that Cx43 was mainly located at the plasma membrane once its expression was restored (Figure 1e). Interestingly, Cx43 was also detected in the nuclear fraction (Figure 1f). Cx43-restored ER+/HER2-BC cells exhibited a restored capacity to communicate through gap junctional intercellular communication (GJIC) when two cells expressing Cx43 get in touch (Figure 1g) and contained active Cx43-hemichannels (HCs) detected by ATP release and under the presence of the (hemi)channel blockers carbenoxolone (CBX) and meclofenamate (Meclo) used a 100 μM and 50 μM respectively for 15 minutes (Figure 1h). Control empty vector (EV) cells exhibited hemichannel activity, likely due to pannexin (Panx1 and Panx2) expression detected by RT-PCR (Figure S1a). However, upregulation of Cx43 further enhanced communication through hemichannels, as illustrated in Figure 1h.

At the functional level, upregulation of Cx43 activity led to a decrease in cellular proliferation, as detected by a colony formation assay in which cells were plated at low density (Figure 1i). A significant reduction in both colony formation capacity and cell proliferation, as evidenced by reduced levels of proliferation marker Ki67 (Figure 1j), was observed after 7 days in culture. These results were validated in a 3D model with ER+/HER2-BC cells (C and Cx43) grown in 3D structures cultured in a U shape plate for 8 days. Cx43 upregulation reduced the area of the 3D structures after 8 days in culture (Figure 1k).

The reduction of cell proliferation and colony formation capacity in 2D and 3D assays correlated with the acquisition of a senescent-like phenotype as showed in Figure 2. Cx43 increased the levels of the senescent marker p21 accompanied by an increase in p53 detected by immunofluorescence (for p21) and western blot (Figure 2a). The β-galactosidase activity assays further confirmed the increase of a senescent phenotype in Cx43-ER+/HER2-cells (Figure 2b). SASP factors were found to be altered following Cx43 restoration, as indicated by changes in mRNA levels (Figure 2c) and in the levels of secreted factors, analyzed using a human proteome profiler array (Figure 2d), when Cx43-restored ER+/HER2-cells were compared to empty vector control cells. Interestingly, Cx43 slightly reduced IL6 involved in BC metastasis or different angiogenic factors, such as PDGF-AA or VEGF. Also, we have found an upregulation of ApoA1 that plays a fundamental tumor-suppressive role and its expression is linked to improved prognosis in certain cancers, highlighting its significance in tumor biology (Figure 2d). The effect of Cx43 in SASPs modulation could be mediated by downregulation of the NFKB signaling as we observed a decrease in NFKB levels when Cx43 is upregulated (Figure 2e). These findings suggest that Cx43 decreases proliferation, enhances cellular senescence, and modulates the expression of secretory factors, leading to a different SASP signature in ER+/HER2-breast cancer cells. However, when we treated Cx43-restored cells with the senolytic drug Navitoclax, our results showed that Navitoclax had little effect on colony-forming capacity, which was already highly decreased in Cx43-restored cells (Figure 2f). However, β-galactosidase activity did not change in wild-type ER+/HER2-breast cancer (BC) cells upon Navitoclax treatment but significantly reduced in Cx43-restored ER+/HER2-cells when treated with Navitoclax (Figure 2g). The colony formation assay (Figure 2f) and β-galactosidase results (Figure 2g) together suggest that although Cx43 enhances senescence (as detected by β-galactosidase activity), additional mechanisms are contributing to its antitumor activity.

**Figure 2.**
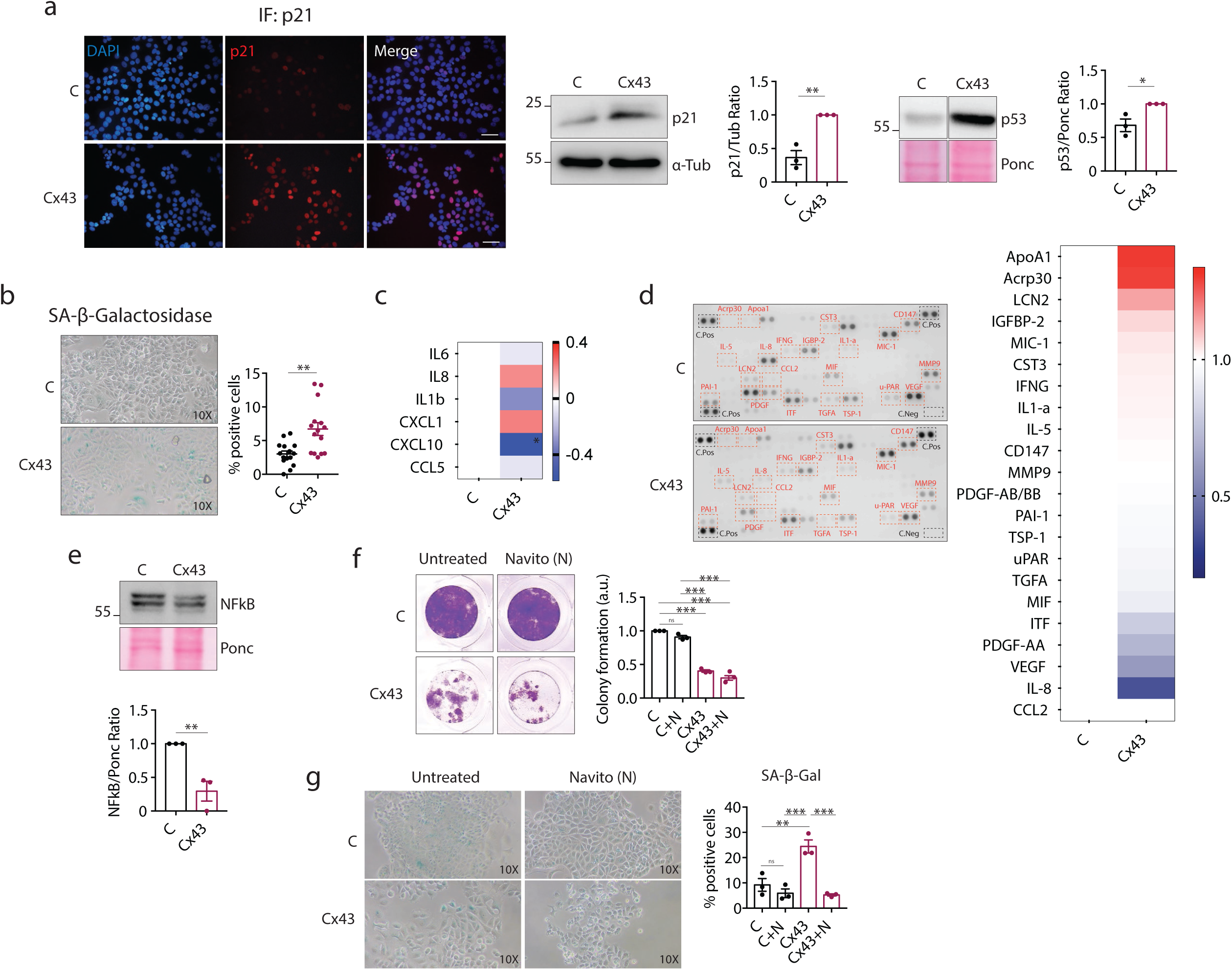
Cx43 induces proliferation arrest and a senescent-like phenotype in ER+/HER2- breast cancer cells. (a) Representative images of p21^CIP1^ (red) immunofluorescence of MCF7 after Cx43 restoration using a lentiviral plasmid. Scale bar: 50 µM. Western blot shows the upregulation of the senescence markers p53 and p21^CIP1^ and a reduction in total RB levels when cells express Cx43. Tubulin and ponceau were used as loading controls. (b) SA-β-galactosidase activity in MCF7 expressing Cx43 shown by light microscopy images. Quantification is shown on the right. Each point on the graph corresponds to the biological replicates within the 3 independent experiments. (c) Heat map of SASP gene expression in MCF7 cells expressing vector or Cx43. The mean of 3 independent experiments is shown. (d) Proteome profile of the conditioned medium from MCF7 C and Cx43 cells. Heat map showing the changes in the SASP components after Cx43 expression (e) Western blot shows the downregulation of NFKB after Cx43 expression. Quantification is shown below. Ponceau was used as loading control. (f) Colony formation assay of MCF7 cells expressing the vector control (C) or Cx43 expression (Cx43) and treated with 3.5μM Navitoclax (N) for 7 days. (g) SA-β-galactosidase activity in MCF7 cells C and Cx43 after treatment with 3.5νM Navitoclax (N) for 7 days shown by light microscopy. The data represent the mean ± SEM of 3-6 independent experiments. Two-tailed Student’s t-test and ANOVA were used to calculate the significance represented as followed: *p<0.05; **p<0.01; ***p<0.001.

### Cx43 modulates the expression of cell cycle mediators and exhibits a cell cycle–enriched interactome when restored in ER+/HER2-breast cancer cells

We next decided to investigate further roles of Cx43 in cell cycle progression and other pathways involved in cell death. Immunofluorescence experiments confirmed a reduction in EdU incorporation on ER+/HER2-BC cells expressing Cx43 compared to the control (Figure 3a). DNA content analysis revealed that Cx43-positive ER+/HER2-BC cells exhibited a prolonged S-phase (and reduced G1 and G2/M) compared to control cells (Figure 3b). Gene expression analysis by qPCR of various cell cycle regulators confirmed a reduction in the expression levels of several key regulators, including CDK4 and CDK6, compared to control (EV) cells, suggesting that Cx43 alters cell-cycle progression leading to an increased in S phase (Figure 3b-c). Since the Cx43 C-terminal domain interacts with several proteins regulating different signalling pathways independently of its channel activity, we performed immunoprecipitation experiments to pull down Cx43 from nuclear (N) and cytoplasmic (C) extracts (Figure 3d) in order to analyse potential interactions with cell cycle regulators. The results revealed that Cx43 interacts with p21 and p27 (as previously reported [39] [40]), both of which are inhibitors of G1 cyclin-CDK protein kinase activity, and that these interactions occur in both the nucleus and in the cytoplasm (Figure 3d).

**Figure 3.**
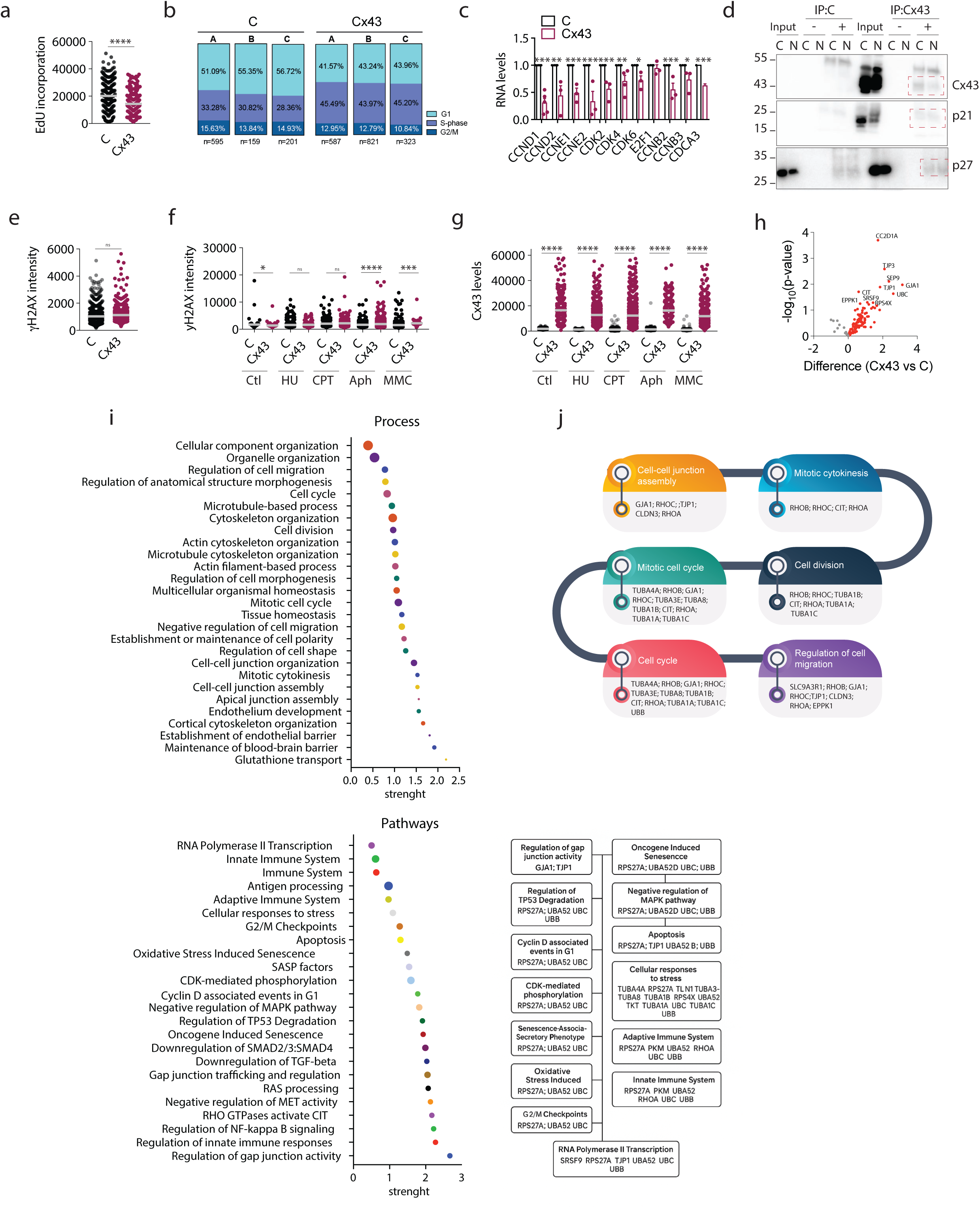
Enhanced Cx43 expression regulates cell-cycle delay in ER+/HER2- BC through the interaction with key cell cycle regulators. (a) Single cell immunofluorescence showing EdU incorporation on MCF7 cells expressing Cx43 compared to the control (C). Each point on the graph corresponds to the biological replicates within the 3 independent experiments. (b) Cell cycle phase distribution in Cx43-positive MCF7 cells compared to the EV control. A, B, C represent 3 independent experiments. (c) Cell cycle regulators expression levels in Cx43-positive cells compared to the control (C). (d) Pull-down assay of Cx43 in nuclear extracts, both in MCF7 control (C) and Cx43-positive cells. Interaction with p27 and p21^CIP1^ was checked. LaminA and Tubulin were used as loading controls. Input corresponds to the whole-cell lysate. IP+ corresponds to the fraction obtained after immunoprecipitation with magnetic beads conjugated to an anti-Cx43 antibody. IP− corresponds to the fraction obtained after incubation with magnetic beads alone and was used as a negative control to assess non-specific binding. (e) Single cell immunofluorescence to check γ-H2AX both in control (C) and Cx43-positive cells. Each point on the graph corresponds to the biological replicates within the 3 independent experiments. (f) Single cell immunofluorescence to check γ-H2AX in both control (C) and Cx43-positive MCF7 cells in combination with different drugs. Each point on the graph corresponds to the biological replicates within the 3 independent experiments. (g) Single cell immunofluorescence to check Cx43 protein levels in Cx43-positive and control cells (C) MCF7 cells in combination with different drugs. Each point on the graph corresponds to the biological replicates within the 3 independent experiments. (h) Volcano plot showing the most abundant proteins identified to interact with Cx43 at the nuclear level. (i) Process and pathway analysis of the proteins identified to interact with Cx43 at the nuclear level. The size of the dots represents the number of proteins identifies in each pathway. Dots size: 2 (0-4 proteins); 4 (4-8 proteins); 6 (8-12 proteins); 7 (12-16 proteins); 9 (16-20 proteins); 10 (>20 proteins). (j) Representative diagram showing the proteins identified to be participated in the cell-cycle related pathways that interact with Cx43. The data represent the mean ± SEM of 3-6 independent experiments. Two-tailed Student’s t-test was used to calculate the significance represented as followed: *p<0.05; **p<0.01; ***p<0.001.

As we have previously observed, Cx43 is also present in the nucleus (Figure 1f), and interacts with nuclear lamins and DNA repair foci [30]. Based on this, we decided to investigate the possibility of accumulated DNA damage under Cx43 restoration in this model as an additional mechanism to explain the observed changes in cell cycle and proliferation. When we examine the levels of the γ-H2AX, a marker of DNA double-strand breaks (DSBs), using single-cell immunofluorescence, we did not find any significant differences in γ-H2AX levels bound at chromatin between control and Cx43-tumor cells (Figure 3e). To further validate these results, we treated cells with different DNA-damaging agents, including hydroxyurea (HU), camptothecin (CPT), aphidicolin (APH), and mitomycin C (MMC) (Figure 3f). Again, no general increase in γ-H2AX levels was detected when Cx43 was upregulated (Figure 3f). However, upregulation of Cx43 itself increased the levels of nuclear Cx43, and treatment with DNA-damaging agents maintained this increase (Figure 3g). These results suggest that DNA damage is unlikely to be responsible for the observed increase in S-phase duration or cell cycle alteration in ER⁺/HER2⁻ BC cells when Cx43 is upregulated. However, Cx43 upregulation appears to exert a chromatin-associated function in ER⁺/HER2⁻ BC cells.

Next, we performed an immunoprecipitation assay of nuclear fractions followed by proteomic analysis to identify proteins interacting with Cx43 in the nucleus. The volcano plot shows the proteins interacting with Cx43 in control and Cx43-tumor cells (Figure 3h and Table 1). Among the interactors, we observed strong enrichment of proteins involved in chromatin remodelling, cell cycle regulation and division, G2/M checkpoints, SASP factor regulation, TP53 degradation, regulation of Cyclin D-associated events during the G1 phase of the cell cycle, programmed oncogene-induced senescence, among others (Figure 3i). These findings indicate that nuclear Cx43 plays broader regulatory roles beyond senescence, cell cycle, or the anchoring of DNA repair foci to the nuclear lamina [30, 38]. We have identified several known Cx43-interactors such as ZO-1 (TJP1) (Figure 3h) [41, 42]. Among key interactors, several tubulin isoforms (TUBA4A, TUBA3E, TUBA8, TUBA1B, TUBA1A, TUBA1C), RHOA, and tight junction protein 1 (ZO-1) were identified within the nucleus, suggesting a role in mitotic processes, cell division, and the regulation of cellular migration and adhesion (Figure 3h-j). Given the effect of Cx43 on the cell cycle (Figure 3a–d), the repertoire of cell-cycle interactors, including proteins related to Cyclin D-associated events and CDK-mediated phosphorylation involving ubiquitin-related proteins such as RPS27A, UBA52, UBC, and UBB, together with the clinical use of cell-cycle inhibitors in ER⁺/HER2⁻ BC, such as CDK4/6 inhibitors (CDK4/6i), we decided to test the activity of Cx43 under CDK4/6 inhibitor treatment.

**Table 1.**
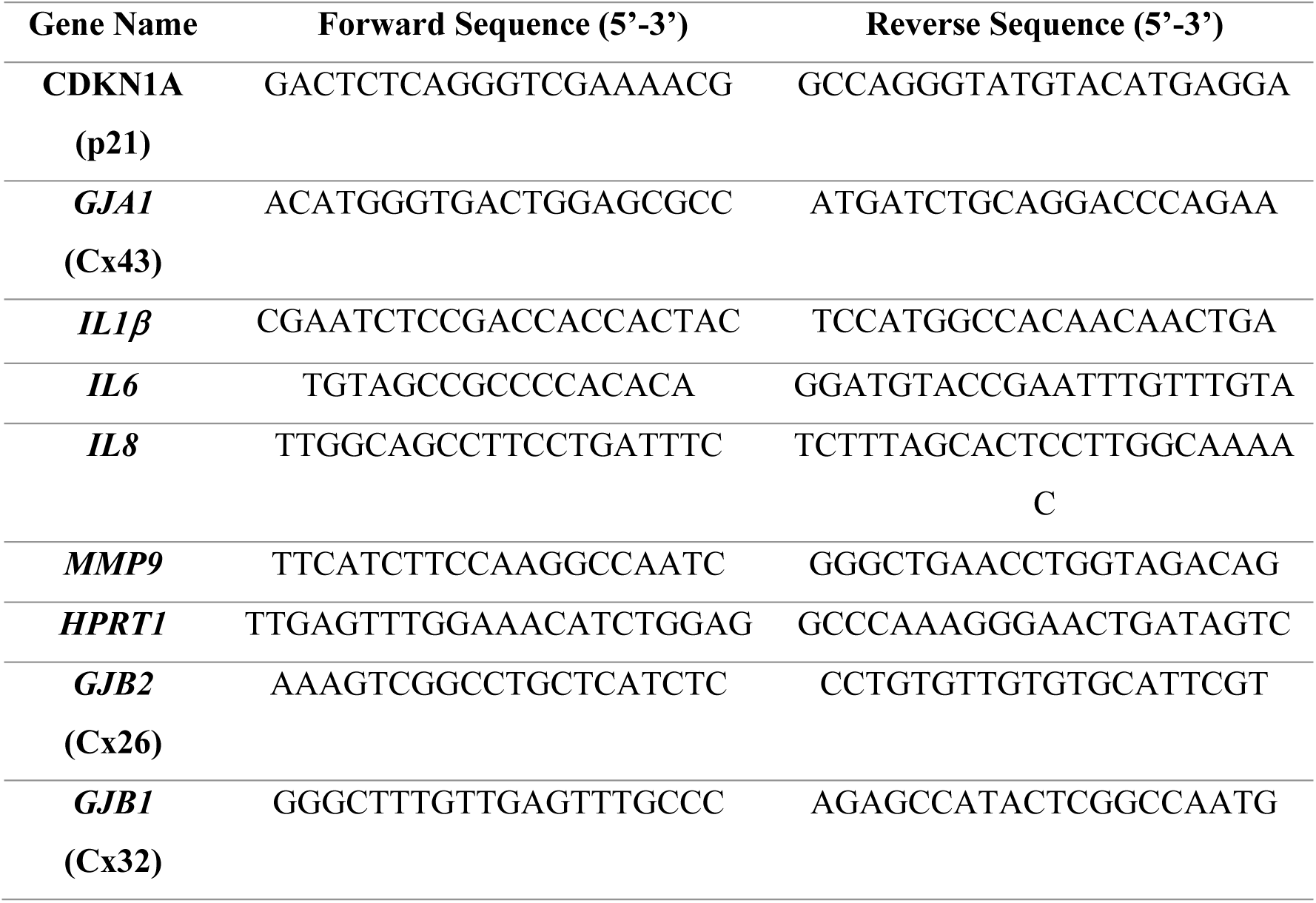
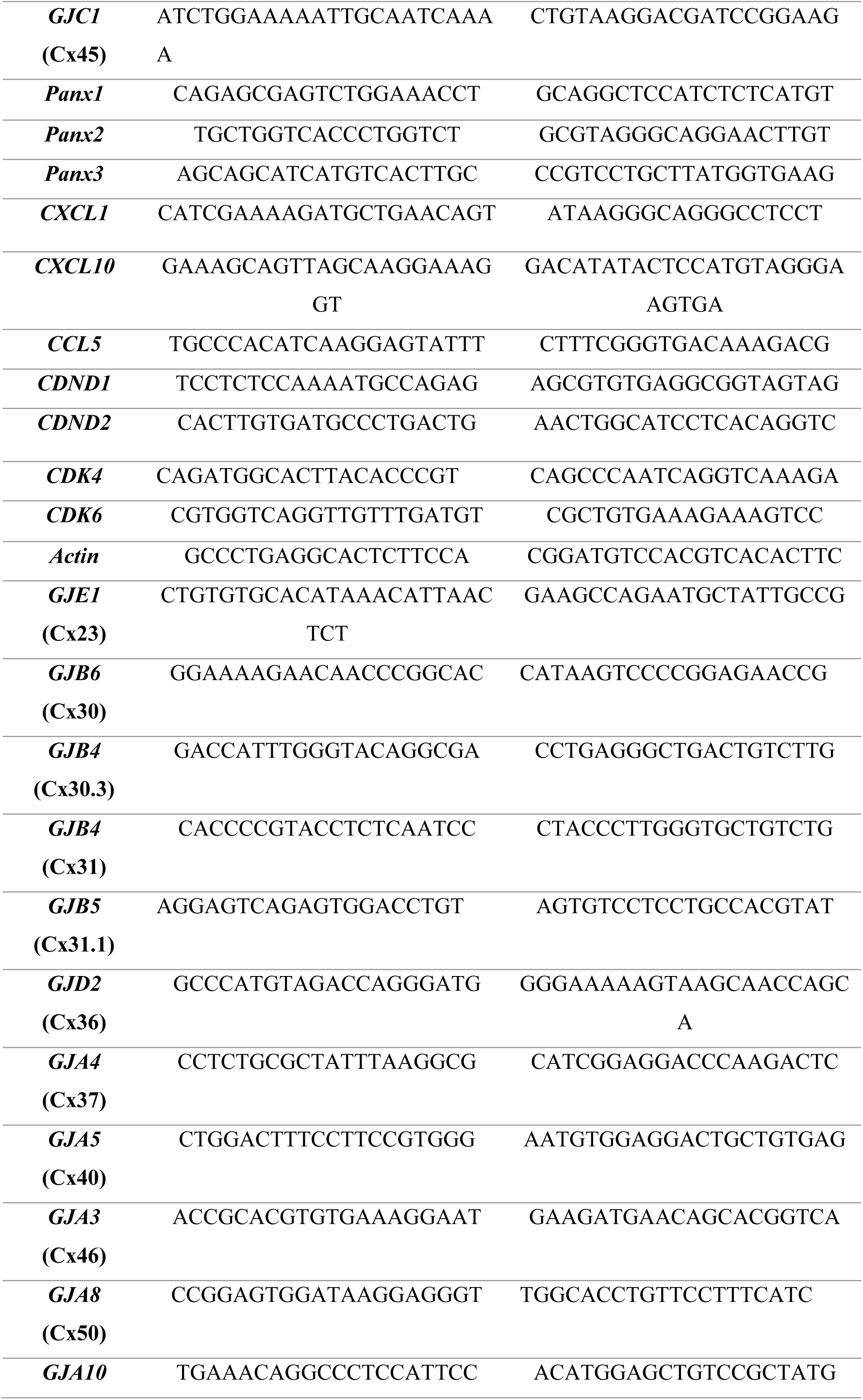
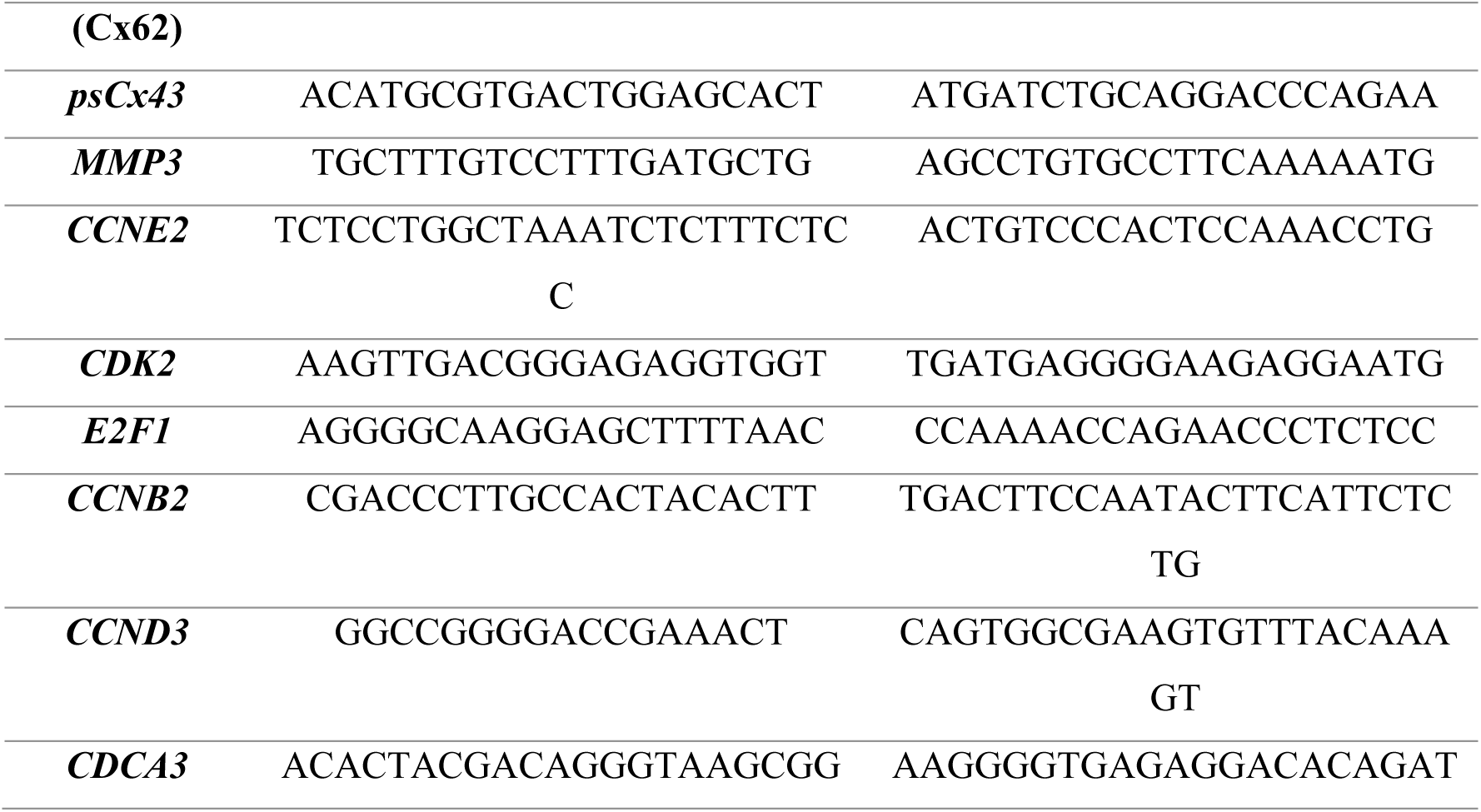
Forward and reverse primer sequences used for gene expression analyses.

### Cx43 Increases the Efficacy of CDK4/6 Inhibitors and the Vulnerability to Senolytic Drugs

Recently, CDK4/6 inhibitors have emerged as a promising therapeutic strategy for cancer treatment, as they can block the G1/S phase transition, inducing proliferation arrest associated with cellular senescence[43]. Based on these results and on the effect of Cx43 on enhancing cellular senescence, we tested the effect of Cx43 restoration under CDK4/6i treatment to determine whether Cx43 reinforces the proliferation arrest and senescent-like phenotype induced by these inhibitors.

MCF7 ER+/HER2-cells (control and Cx43-positive cells) were treated with 250nM Palbociclib for 7 days (Figure 4a). Western blot and immunofluorescence assays revealed that Cx43 levels increased during Palbociclib treatment, particularly in Cx43-MCF7 cells, consistent with the treatment response (Figure 4a–b). qPCR analysis after Palbo treatment revealed increased mRNA levels of Cx43, Cx32, and Cx46 (Figure S2a. Next, both ER+/HER2-BC cells expressing the vector control (C) and Cx43 were treated with different concentrations of Palbociclib and the results showed that Cx43 reduced by 4 times the IC50 of CDK4/6i from 282,1 nM (C cells. IC₅₀ 95% CI: 82.23 -1393) to 66,82 nM (Cx43 cells. IC₅₀ 95% CI: 0.01618 -240,805) (Figure 4b). As Cx43 levels vary among luminal breast cancer cells (Figure S2b), we included a different ER+/HER2− BC cell line with higher Cx43 expression (T47D; Figure 4c) that effectively respond to CDK4/6i (Figure S2c) and observed that, in ER+/HER2− BC cells, Cx43 levels are directly correlated with an increased response to CDK4/6i (Figure 4d-e). The ER+/HER2− BC cell line T47D expresses endogenously lower levels of Cx43 compared to MCF7-Cx43 cells, but higher than MCF7 control cells (C), as detected by Western blot and IF (Figure 4c; Figure S2d). T47D cells exhibited a lower proliferation rate (Figure S2e) and higher β-galactosidase activity (Figure S2f) compared to MCF7 cells at the basal level. When treated with increasing concentrations of the CDK4/6 inhibitor Palbociclib, T47D cells showed a stronger response to the drug than MCF7 cells, with an IC50 of 120.6 (Figure 4d–e). Together, these results indicate that Cx43 protein levels correlate with Palbociclib response in tumor cells (MCF7 and T47D). Specifically, higher levels of Cx43 were directly associated with a lower Palbociclib IC50 (Figure 4e), confirming confirming that Cx43 restoration enhances the efficacy of the CDK4/6 inhibitor palbociclib.

**Figure 4.**
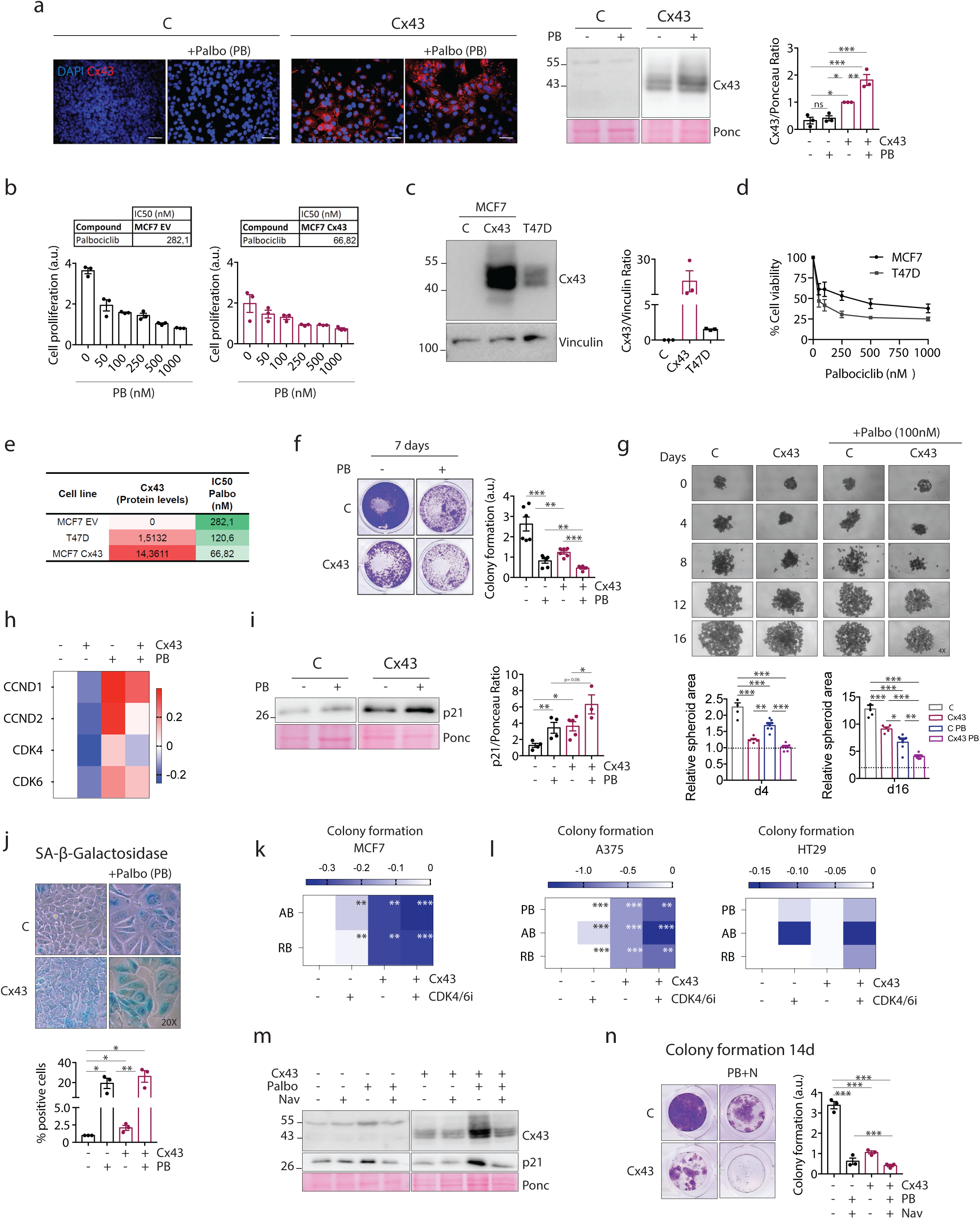
Cx43 sensitizes ER+/HER2 breast cancer to CDK4/6 inhibitors increasing their efficacy. (a) Immunofluorescence to confirm Cx43 increased expression and cell location after treating MCF7 EV and Cx43 with PB 500 nM for 7 days. Scale bar: 50 µM. Western blot for Cx43 protein levels after 7 days of treatment with 500nM Palbociclib (PB) compared to the untreated control. The graph represents the quantification of 3 independent experiments. Ponceau was used as loading control. (b) Palbociclib dose-response in MCF7 C and Cx43-positive cells. IC50 calculation is shown on the tables. (c) Western-blot comparing Cx43 protein levels among control (C) and Cx43-positive MCF7 and T47D breast cancer cell lines. Vinculin was used as loading control. Quantification is showed on the right. (d) Cell viability of MCF7 and T47D cells measured by crystal violet staining (A_570_). (e) Table showing the correlation between Cx43 protein expression and CDK4/6i Palbociclib through its IC50 in MCF7 control (C) and Cx43-positive and T47D cells. (f) Colony formation capacity of MCF7 control and Cx43 cells after 7 days treatment with Palbociclib 250 nM. (g) Spheroids of MCF7 control (C) and Cx43-positive cells treated with PB 100nM for 16 days. Quantifications are shown below. (h) RNA expression levels of CDK4/6i targets in MCF7 C and Cx43 treated with PB 250nM for 7 days. (i) p21^CIP1^protein levels in MCF7 after Cx43 ectopic expression and 7 days Palbo treatment. Ponceau was used as loading control. (j) SA-β-galactosidase activity in MCF7 expressing Cx43 after 7 days of Palbo treatment, shown by light microscopy images. (k) Heatmap representing the colony formation capacity of MCF7 cells after treatment with Abemaciclib (1μM) and Ribociclib (500nM) for 7 days. (l) Heatmap representing the colony formation capacity of A375 and HT29 cells after treatment with Palbociclib (500nM), Abemaciclib (1μM) and Ribociclib (500nM) for 7 days. (m) Cx43 and p21^CIP1^protein levels in MCF7 after Cx43 ectopic expression, 14 days Palbo treatment and 7 days of Navitoclax treatment. Ponceau was used as loading control. (n) Colony formation assay of MCF7 cells expressing the vector control (C) or Cx43 ectopic expression (Cx43) and treated with 500nM Palbociclib for 14 days and 3.5μM Navitoclax (N) for 7 days. Quantification for the triple combination is shown on the right The data represent the mean ± SEM of 3-6 independent experiments. Two-tailed Student’s t-test and ANOVA were used to calculate the significance represented as followed: *p<0.05; **p<0.01; ***p<0.001.

To further demonstrate the enhanced antitumor effect resulting from the combination of Cx43 and Palbociclib, ER+/HER2− BC (MCF7) cells were treated for 7 days with Palbociclib, and cell proliferation was assessed using a colony formation assay. The combination of Cx43 and Palbociclib significantly reduced the colony-forming capacity of ER+/HER2− BC cells in 2D (Figure 4f) and in a 3D spheroid model using 100 and 250 nM of Palbociclib concentrations (Figure 4g; Figure S2g). Changes in key cell cycle regulators involved in CDK4/6i response (CCND1, CCND2, CDK4, CDK6) were analysed by qPCR (Figure 4h). To determine whether Cx43 reinforces the senescence-like phenotype induced by Palbociclib in BC cells, we analyzed the senescence markers p21 and β-galactosidase activity (Figure 4i–j). The data showed that the combination of Cx43 and Palbociclib increased both p21 expression (Figure 4i) and β-galactosidase activity (Figure 4j) in ER+/HER2− BC cells after 7 days of treatment. The enhanced effect was also observed with the other CDK4/6 inhibitors, Abemaciclib and Ribociclib (Figure 4k), as well as in different tumor types, including A375 human melanoma and HT-29 human colon carcinoma, which are responsive to CDK4/6 inhibition (Figure 4l). Vector-mediated upregulation of Cx43 significantly enhanced the efficacy of Palbociclib, Abemaciclib, and Ribociclib, as shown by a marked reduction in colony-forming capacity after 7 days of treatment (Figure 4l).

We next assessed whether Cx43-enhanced senescence induced by CDK4/6 inhibition would generate a phenotype sensitive to the senolytic agent Navitoclax (ABT263). The triple combination of Cx43 upregulation, 250 nM Palbociclib, and 3.5 μM Navitoclax significantly reduced colony formation and decreased the expression of the senescence marker p21, representing the most effective strategy to eliminate ER+/HER2− tumor senescent cells after 14 days of treatment (Figure 4m–n). The reduction of Cx43 and p21 expression likely reflects the loss of Cx43-positive senescent cells under Navitoclax treatment after 14 days treatment (Figure 4m).

Next, to further investigate the mechanisms underlying the enhanced response to CDK4/6 inhibition in Cx43-expressing ER+/HER2− breast cancer (BC) cells, we characterized the Cx43-associated protein network under untreated conditions and following palbociclib treatment (Figure 5). For this purpose, we performed comparative mass spectrometry analysis of Cx43 immunoprecipitates (IP+) obtained from total cell lysates of Cx43-positive cells treated with palbociclib (PB) for 7 days and untreated controls (UT) (Figure 5 and Figure S2h).

**Figure 5.**
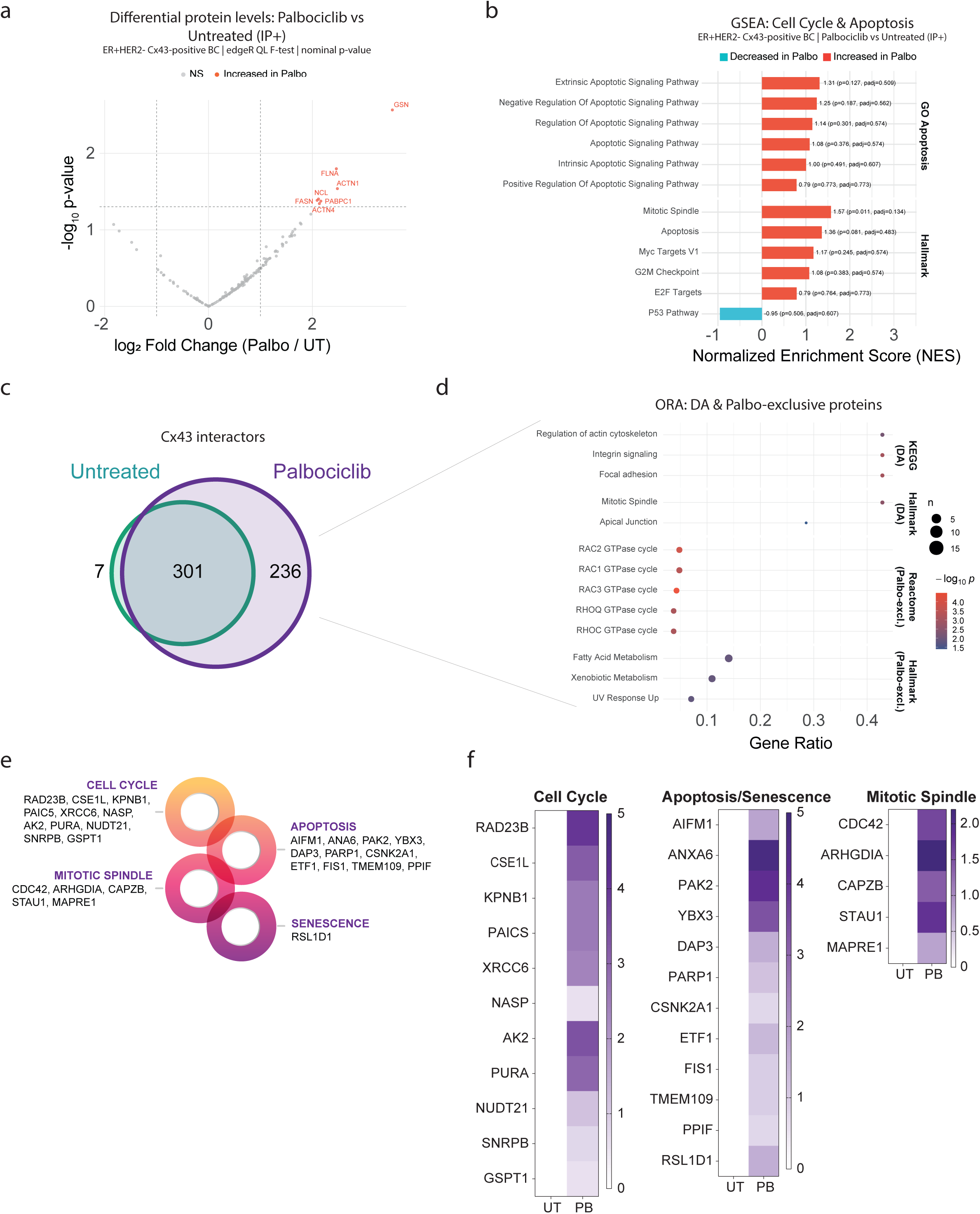
Proteomic analysis of Cx43-immunoprecipitates from Cx43-expressing ER+/HER2- BC cells after Palbociclib treatment. (a) Volcano plot showing differential protein abundance analysis performed using edgeR (quasi-likelihood F-test) on raw spectral counts with trimmed mean of M-values (TMM) normalization in Cx43-expressing ER+/HER2− breast cancer cells under untreated conditions (UT) or following 7 days of palbociclib treatment (Palbo). The plot displays log_2_ fold change (Palbo/UT) versus −log_10_ nominal p-value. The 7 differentially abundant proteins (nominal p < 0.05, |log2FC| > 1) are highlighted in red. All differentially abundant proteins exhibited increased abundance following palbociclib treatment. (b) Gene Set Enrichment Analysis (GSEA) bar plot showing enrichment of selected Gene Ontology (GO) and Hallmark gene sets associated with cell cycle regulation and apoptosis. Bars represent the normalized enrichment score (NES), with nominal p-values and the number of genes (n) indicated for each gene set. Gene sets are grouped according to their respective functional categories. (c) Venn diagram showing the distribution of proteins: 236 proteins were exclusively identified in the Palbo group (detected in ≥2 samples), 7 proteins were exclusive to the UT group, while 301 proteins were shared between both conditions. (d) Combined Over-Representation Analysis (ORA) dot plot of differentially abundant (DA) proteins and Palbo-exclusive (Palbo-excl.) proteins. Enrichment analysis of the 7 differentially abundant proteins using KEGG and Hallmark gene sets identified pathways associated with focal adhesion, integrin signaling, and cytoskeletal organization. Analysis of the 236 Palbo-exclusive proteins using Reactome and Hallmark gene sets revealed enrichment of Rho GTPase-related pathways, including RAC2 and RAC3 GTPase cycles, as well as lipid metabolism-related processes. Dot size represents the number of proteins associated with each pathway (n), dot color indicates −log10 p-value, and the x-axis represents the gene ratio. (e) Schematic representation of selected Cx43-interacting proteins associated with cell cycle regulation, apoptosis, mitotic spindle organization, and senescence-related pathways following palbociclib treatment. (f) Heatmaps showing the spectral counts of selected proteins associated with cell cycle regulation, apoptosis/senescence, and mitotic spindle organization.

Differential protein abundance analysis revealed a subset of proteins enriched in Cx43 immunoprecipitates following palbociclib treatment (p < 0.05, |log2FC| > 1). Interestingly, all proteins meeting these criteria exhibited increased abundance in the palbociclib-treated condition, suggesting treatment-associated changes in the composition of the Cx43-associated protein network (Figure 5a). To further investigate the biological processes associated with these changes, Gene Set Enrichment Analysis (GSEA) was performed, focusing on pathways related to cell cycle regulation and apoptosis. This analysis revealed positive normalized enrichment scores (NES) of several Gene Ontology (GO) terms associated with apoptotic signaling, including intrinsic and extrinsic apoptotic signaling pathways and the positive and negative regulation of apoptosis (Figure 5b). In addition, Hallmark gene sets associated with cell cycle regulation, including *Mitotic Spindle*, *G2M Checkpoint*, *E2F Targets*, and *MYC Targets V1*, exhibited positive NES, whereas the *P53 Pathway* showed negative enrichment. These findings indicate that palbociclib treatment is associated with changes in the representation of Cx43-associated proteins involved in cell cycle control and apoptotic signaling.

Comparative proteomic profiling identified 236 proteins exclusively detected in Cx43 immunoprecipitates following palbociclib treatment, compared with only 7 proteins exclusive to untreated cells, while 301 proteins were shared between both conditions (Figure 5c). These results revealed substantial differences in the composition of the detectable Cx43-associated proteome following CDK4/6 inhibition. Functional enrichment analysis was subsequently performed using Over-Representation Analysis (ORA) to characterize the biological processes associated with differentially abundant and palbociclib-exclusive proteins (Figure 5d). Analysis of differentially abundant proteins identified enrichment of pathways related to the regulation of the actin cytoskeleton, integrin signaling, and focal adhesion, together with Hallmark gene sets associated with mitotic spindle organization and apical junctions. Interestingly, analysis of the 236 palbociclib-exclusive proteins revealed enrichment of several Rho GTPase-related pathways, including RAC1, RAC2, RAC3, RHOQ, and RHOC GTPase cycles. Additional enrichment was observed for Hallmark gene sets related to fatty acid metabolism, xenobiotic metabolism, and ultraviolet response. Collectively, these findings suggest that palbociclib treatment is associated with changes in the Cx43-associated protein network involving cytoskeletal organization, cell adhesion, and intracellular signaling pathways.

Given the involvement of Cx43 in cell cycle regulation and the enhanced response to CDK4/6 inhibition, we next focused on Cx43-associated proteins implicated in cell cycle progression, apoptosis, mitotic spindle organization, and senescence (Figure 5e). Among these, several proteins involved in cell cycle-related processes were identified, including RAD23B, CSE1L, KPNB1, PAICS, XRCC6, NASP, AK2, PURA, NUDT21, SNRPB, and GSPT1. In addition, proteins associated with apoptotic processes, such as AIFM1, ANXA6, PAK2, YBX3, DAP3, PARP1, CSNK2A1, ETF1, FIS1, TMEM109, and PPIF, were detected. Furthermore, proteins involved in mitotic spindle organization and cytoskeletal dynamics, including CDC42, ARHGDIA, CAPZB, STAU1, and MAPRE1, were identified, together with RSL1D1, a protein associated with senescence-related processes (Figure 5e-f). In particular, proteins associated with cell cycle regulation, apoptosis, and mitotic spindle organization displayed distinct abundance patterns between conditions, further supporting treatment-associated changes in the Cx43-associated proteome. Taken together, these proteomic findings indicate that palbociclib treatment is associated with substantial remodeling of the Cx43-associated protein network, particularly involving proteins related to cell cycle regulation, apoptotic signaling, cytoskeletal organization, and senescence-associated processes. These observations support a potential role for Cx43 as a signaling hub connecting multiple cellular processes and suggest that changes in its associated protein network may contribute to the enhanced response to CDK4/6 inhibition observed in Cx43-expressing ER+/HER2− BC cells.

### Nanovesicle-Mediated mRNA Delivery of Cx43 Potentiates the Therapeutic Effect of CDK4/6 Inhibitors

We next decided to test a therapeutic strategy distinct from gene-based therapy, aimed at restoring Cx43 expression in tumor cells. Based on our expertise in using extracellular vesicles (sEVs) as carriers for delivering therapeutic proteins and mRNA [30], we employed these nanovesicles for the delivery of Cx43 into target tumor cells. These sEVs can be produced either from genetically modified tumor cells or from healthy non-transformed cells, such as HEK293T. First, we restored Cx43 expression in ER+/HER2− BC cells using retroviral vectors (Figure 1e), which led to the production of sEVs enriched in Cx43 protein (Figure S3a-c). Cx43 upregulation did not alter the number of secreted particles per cell or the size of EVs, as determined by Nanoparticle Tracking Analysis (NTA) (Figure S3b). However, we observed an increase in their internalization efficiency, as determined by IF detection of nanovesicles labeled with DilC (Figure S3c), compared to sEVs derived from control empty vector-transduced cells. When ER+/HER2-breast cancer (BC) cells were treated with Cx43 enriched sEVs we observed a reduction in the colony formation capacity compared to tumor cells treated with EVs isolated from the EV control (Figure 6a). These results suggest that modification of ER+/HER2− BC cells to express Cx43 leads to the release of particles with antitumor activity, thereby amplifying the antitumor effect of Cx43. Next, to obtain EVs from a non-tumor source already validated in clinical trials for engineered sEVs production (Clinical Trial NCT04592484, registered on October 19, 2020) [44], we used the embryonic cell line HEK293T cells. HEK293T sEVs endogenously express Cx43 (Figure 6b). sEVs were characterized using NTA (Figure 6c) and the internalization of sEVs in MCF7 target cells was analysed by labelling with DilC for 30 minutes and incubated with tumor cells overnight and detected by fluorescent microscopy (Figure 6d). Isolated Cx43-enriched sEVs were used to treat two different ER+/HER2− BC cell lines (Figure 6e). As shown, co-treatment of MCF7 and T47D cells with Cx43+ sEVs and CDK4/6 inhibitors for 7 days reduced colony-forming capacity and increased the efficacy of CDK4/6 inhibition. Using this approach, we confirmed the effect of Cx43 in ER+/HER2− BC cells and how Cx43 potentiates CDK4/6 inhibition and established a strategy for delivering Cx43 mRNA into tumor cells to enhance CDK4/6i efficacy.

**Figure 6.**
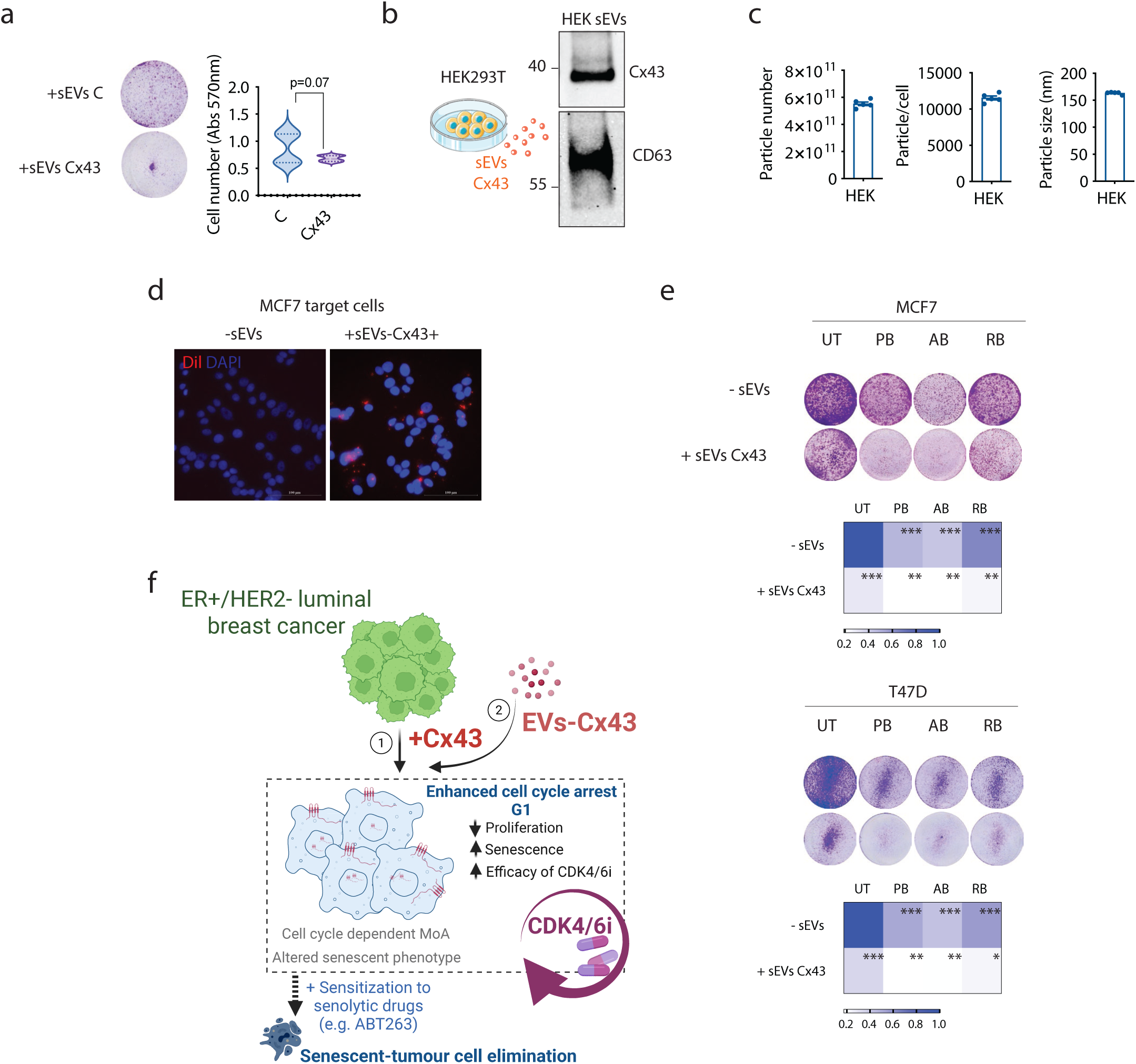
sEVs containing high levels of Cx43 have anti-tumor activity and improve CDK4/6i efficacy. (a) Colony formation assat of MCF7 cells treated with sEVs isolated from EV (sEVs C) and Cx43 (sEVS Cx43) for 7 days. SeVs were refreshed every 48 hours. (b) Western-blot showing that sEVs isolated from HEK 293 contain high levels of Cx43. CD63 was used as marker to confirm the presence of sEVs. (c) Nanoparticle Tracking Analysis (NTA) of sEVs isolated from HEK293T. Graphs represent the particle number, particles/cell and particles size. (d) Overnight internalization experiment of HEK isolated sEVs in MCF7 target cells. -sEVs (untreated controls), sEVs-Cx43+ (sEVs derived from HEK-293). Scale bar: 100 µM. (e) Colony formation assay of MCF7 and T47D breast cancer cells treated with 500nM of Palbociclib, 1μM of Abemaciclib and 1μM of Ribociclib alone or in combination of sEVs positive for Cx43 isolated from HEK293 for 7 days. The presence of sEVs enriched in Cx43 decreases the capacity to form colonies and increases the efficacy of treatment in both cell lines. Treatments and sEVs were refreshed every 48 h. -sEVs (untreated controls), sEVs-Cx43+ (sEVs derived from HEK-293). (f) Graphical abstract: restoration of Cx43 in ER+/HER2− BC cells reduce cell proliferation and promotes the activation of a distinct senescence phenotype. The combination of Cx43 and CDK4/6 inhibitor, enhances inhibitors efficacy through cell cycle–dependent mechanisms and rendering cells more susceptible to senolytic agents such as Navitoclax. The data represent the mean ± SEM of 3 independent experiments. Two-tailed Student’s t-test was used to calculate the significance represented as followed: *p<0.05; **p<0.01; ***p<0.001.

## DISCUSSION

In this study, we explored the tumor-suppressive properties of Cx43 in ER+/HER2− breast cancer. Our findings demonstrate that restoration of Cx43 in ER+/HER2− BC cells reduce cell proliferation and promotes the activation of a distinct senescence phenotype. Importantly, the combination of Cx43 and CDK4/6 inhibitors emerged as the most effective strategy to suppress tumor cell proliferation in this BC subtype, enhancing CDK4/6i efficacy through cell cycle– dependent mechanisms and rendering cells more susceptible to senolytic agents such as Navitoclax.

Recent studies have highlighted the importance of Cx43 as a breast tumor suppressor [22]. However, this role appears to depend on the breast cancer subtype [22, 33]. Our results show that Cx43 is diffusely expressed in tumor samples from patients with ER+/HER2− BC (Figure 1a–b), and database analyses (Metabric) confirm that Cx43 upregulation is a favourable prognostic factor. High Cx43 levels were associated with improved overall survival probability (Figure 1c). However, while a significant association was observed in the METABRIC cohort, the TCGA and Lu datasets did not reach statistical significance, and the Wang dataset showed a different trend (Figure S1b), emphasizing that the prognostic value of Cx43 expression may be cohort-dependent and requires further validation in larger patient series. In addition, it should be noted that Cx43 mRNA levels do not necessarily correlate with protein abundance, localization, or functional activity, as Cx43 has a rapid protein turnover, and its functionality is subject to multiple post-transcriptional and post-translational regulatory mechanisms. Therefore, transcriptomic datasets may not fully capture the biological effects associated with Cx43 protein expression. Importantly, restoration of Cx43 in ER+/HER2− BC models reduced colony formation capacity and proliferation in both 2D (Figure 1i–j) and 3D models (Figure 1k). Together, our findings not only reveal the tumor-suppressive role of Cx43 in the ER+/HER2− BC subtype but also underscore its potential as a prognostic biomarker.

Gap-junction intercellular communication (GJIC) is often lost in various human tumors compared with adjacent normal tissue [23]. Restoration of GJIC in tumor cells, particularly at early stages of the disease, has been shown to provide therapeutic benefits in several in vitro and in vivo models [45, 46]. In this study, we demonstrated that Cx43 localizes to the plasma membrane after its restitution in BC cells, thereby restoring both GJIC and hemichannel activity (Figure 1e–j). Interestingly, our results also reveal that Cx43 is present at cytoplasmic and nuclear levels, where it exerts direct effects on cell cycle regulation (Figure 3). These findings indicate that the role of Cx43 in BC cells extends beyond its canonical channel functions. Although our data suggest that Cx43 may act in a channel-independent manner, further studies are required to determine whether its nuclear activity is functionally linked to, or independent from, its channel activity [30, 47].

Previous reports in ovarian carcinoma, osteosarcoma, glioma, and other models have shown that upregulation or restoration of Cx43 inhibits cell proliferation by inducing G1 cell cycle arrest, often through increasing levels of p27 (a Cip/Kip family member), sometimes by reducing Skp2-mediated degradation of p27 or by other regulatory mechanisms, or by delaying in mitotic progression [29, 40]. Other studies have demonstrated that Cx43 interacts with Cyclin E and plays a critical role in regulating the G1 to S phase transition of the cell cycle [48]. Our study identified a cell cycle alteration in Cx43-expressing cells, characterized by an accumulation of cells in S-phase and an increase in mitotic duration (Figure 3a–d). We also observed reduced expression of key cell cycle–related genes (e.g., CCND1, CDK2, CDK4, E2F) (Figure 3c) in Cx43-positive cells. Furthermore, proteomic analyses revealed that Cx43 interacts at the nuclear level with proteins such as RPS27A, UBA52, UBC, and UBB (Figure 3i), whose expression is tightly regulated during the cell cycle, particularly in S-phase and G2/M. This suggests a potential role for Cx43 in ribosome biogenesis and proteostasis during rapid cell division, thereby linking Cx43 to broader changes in cell cycle control (Figure 3). Collectively, these results support the idea that Cx43 exerts tumor-suppressive effects by modulating the balance between positive and negative regulators of cell cycle progression. Importantly, unlike previous studies reporting that Cx43 acts solely by reducing proliferation [48], our findings highlight a broader role that also includes the induction of cellular senescence and modulation of SASP expression.

The induction of a senescent-like phenotype in cancer cells has emerged as a useful therapeutic strategy, initially with chemo-and radiotherapy and more recently with specific drugs that target senescence [49]. The efficacy of CDK4/6 inhibitors has been studied in ER+ BC subtype [50–54], and they are known to act by inducing tumor cell senescence [55–57]. However, their precise mechanisms of action remain unclear, and although certain vulnerabilities have been described [20], no molecular targets have yet been identified that can reliably enhance their efficacy. Our findings suggest that sensitivity to the inhibitors may be associated with Cx43 expression levels in ER+/HER2− BC models. However, given the differences in the genetic backgrounds of the cell lines included in this study, further validation is required to confirm this association. Importantly, senescence plays multiple roles in health and disease: while acute senescence is beneficial—for example, by limiting fibrosis [58] and cancer initiation [59]—persistent senescence can be detrimental [60, 61]. Although CDK4/6i are effective at halting cancer cell proliferation, accumulating evidence indicates that therapy-induced senescent cells may persist in tissues, becoming resistant to cell death and often refractory to senolytic drugs [20]. These tumor senescent cells, together with SASP factor induction, promote inflammation, tissue dysfunction, and may support tumor progression or relapse. For instance, although Abemaciclib provides clinical benefit, the long-term presence of therapy-induced senescent cells contributes to disease recurrence and reduced treatment efficacy over time [20]. Therefore, the use of senolytics—drugs that selectively eliminate senescent cells—has been proposed as a complementary strategy to improve treatment outcomes [62]. Our results demonstrate that restoration of Cx43 in BC cells not only enhances the efficacy of Palbociclib and other CDK4/6i, but also renders these tumor senescent cells susceptible to the senolytic drug Navitoclax.

The SASP-mediated paracrine effects released by senescent cells can remodel the surrounding tissue by modulating the behaviour of adjacent cells [30, 31, 63]. Among these SASP factors, IL-6, whose expression is reduced upon Cx43 restoration, appears to have prognostic significance. When active, IL-6 can promote BC metastasis and suppress the anti-tumor immune response [32, 33]. We also detected downregulation of angiogenic factors such as PDGF-AA and VEGF upon Cx43 upregulation. These findings suggest that Cx43 not only exerts antitumor activity in ER+/HER2− BC cells by affecting the cell cycle and inducing senescence but also shifts the senescence phenotype toward a more favourable SASP profile, reducing factors that drive disease progression.

Previous studies have implicated Cx43 in different aspects of tumor suppression, including contact inhibition and modulation of signalling pathways such as p53, mTOR, and NF-κB [22, 64]. Our immunoprecipitation results revealed substantial changes in the Cx43-associated protein network following Palbociclib treatment, identifying 236 proteins exclusively detected in treated cells, compared with only 7 in untreated controls, while 301 proteins were shared between both conditions (Figure 5). Interestingly, functional enrichment analysis revealed an increased representation of proteins involved in cell cycle regulation, mitotic spindle organization, and apoptotic signalling following Palbociclib exposure. Particularly, several proteins associated with cell cycle control, including RAD23B, CSE1L, KPNB1, PAICS, and XRCC6, were identified among Cx43-associated proteins (Figure 5), suggesting a potential involvement of Cx43 in the regulation of cell cycle-related processes during CDK4/6 inhibition. In addition, proteins implicated in apoptotic signalling, such as AIFM1, ANXA6, PAK2, YBX3, and PARP1, were also detected, supporting a possible connection between Cx43-associated protein networks and cell death-related mechanisms. Furthermore, enrichment of pathways related to actin cytoskeleton regulation, focal adhesion, and Rho GTPase signalling, particularly RAC1, RAC2, RAC3, RHOQ, and RHOC, suggests that Palbociclib treatment may also influence Cx43-associated processes involved in cytoskeletal dynamics and cell adhesion. These processes are closely connected to cell cycle progression and cellular responses to antiproliferative stimuli [65]. Besides, the identification of proteins associated with mitotic spindle organization, including CDC42, ARHGDIA, CAPZB, STAU1, and MAPRE1, together with the senescence-associated protein RSL1D1, further highlights the involvement of the Cx43-associated proteome in pathways related to proliferative control and cellular senescence [66]. Collectively, these findings suggest that Cx43 may function as a signalling hub connecting cell cycle regulation, apoptotic signalling, and cytoskeletal organization, potentially contributing to the enhanced response to CDK4/6 inhibition observed in Cx43-expressing ER+/HER2− breast cancer cells.

We also explored an alternative strategy to restore Cx43 in BC cells by using nanovesicles for targeted delivery. Extracellular vesicles (EVs) are considered one of the most promising nano-delivery systems, and Cx43 has been shown to be a key factor for efficient cargo transfer to recipient cells a finding further supported by our data (Figure S3a). The use of EVs as therapeutic nano-carriers offers several advantages, including high biocompatibility, low toxicity, and low immunogenicity for the delivery of proteins and mRNA [67, 68] . Our data demonstrate that EVs enriched in Cx43, isolated from both tumoral and non-tumoral cell lines, reduced colony-forming capacity and significantly increased CDK4/6i efficacy in in vitro BC models. These findings highlight EV-based delivery as a promising approach for restoring Cx43 activity in tumors, consistent with our previous observations in other cancer types [30]. However, it remains unclear whether the observed effect is due to direct protein transfer or to transcriptional changes induced by the treatment of target cells with Cx43-enriched EVs.

In summary, our study demonstrates that Cx43 exerts an anti-proliferative effect in ER+/HER2− BC cells and, in combination with CDK4/6 inhibitors, enhances therapeutic efficacy. Furthermore, Cx43-dependent senescence and the associated changes in SASP composition enhance sensitivity to senolytic drugs such as Navitoclax. We also propose a novel therapeutic strategy based on EV-mediated delivery of Cx43 to ER+/HER2− BC models (Figure 6f). Importantly, these findings may be extended to other tumor types, with potential clinical applications to improve the overall efficacy of CDK4/6i-based therapies and enhance patient outcomes.

## MATERIALS AND METHODS

### Cell culture

Human primary tumour cells were grown following previously published protocols [69]. Human cell lines MCF7 (breast cancer), T47D (breast cancer) and A375 (melanoma) were cultured in Dulbeccós modified Eaglés medium (DMEM) (Lonza) supplemented with 10 % fetal bovine serum (FBS) (Gibco, Thermo Fisher Scientific) and 100 U/mL penicillin and 100 μg/ml streptomycin (Gibco, Thermo Fisher Scientific). HT29 (colon adenocarcinoma) was cultured in McCoy’S 5A medium (Sigma-Aldrich) supplemented with 10 % FBS and 100 U/mL penicillin and 100 μg/ml streptomycin. Cells were maintained at 37 °C in 5 % CO_2_ humidified incubator (SANYO CO^2^). The medium was changed every 2 days.

### Retroviral infection

Retroviral infection was used to ectopically express Cx43 in MCF7 cells. Retroviral particles were generated by transfecting pLPCX-Cx43 and pLPCX-EV plasmids, kindly donated by Dr. Trond Aasen, and retroviral helper plasmids (vsvg and gag-pol) with Polyethylenimine (PEI) in HEK293T packaging cells for 48h. The supernatant was then filtered with 0.45µm filters and applied to MCF7 cells in the presence of 4µg/ml polybrene (hexadimethrine bromide; Sigma-Aldrich) following 3 rounds of infection. Cells were subsequently selected with the appropriate antibiotic resistance using 0.5μg/ml puromycin (Sigma).

### Cell Transfection

Cell lines (A375, HT29 and T47D) were transfected by electroporation to ectopically express Cx43. The Amaxa® Cell Line Nucleofector® Kit V (Lonza) was used to transfect cells in a Cell Line Nucleofector™ Device (Lonza). A million cells were harvested and resuspended in 100 μL of Nucleofector® solution. 3 μg of the corresponding plasmid was added, and the mixture was transferred into an electroporation cuvette using the U-28 program. Cell lines were transfected with pIRESpuro2 plasmid construct (Clontech) containing the human Cx43 sequence, kindly donated by Arantxa Tabernero (Institute of Neuroscience of Castilla y León, University of Salamanca, Spain). The transfected cells were seeded in culture medium and 24 hours (h) post-transfection the medium was replaced with complete medium containing puromycin dihydrochloride (Tocris, Bioscience) at different concentrations depending of the cell line (0,5 μg/mL to 2 μg/mL).

### CDK4/6 inhibitors treatment

Cells were treated with 250nM or 500nM of Palbociclib (PD0332991) (APExBIO), 500nM Abemaciclib (LY2835219) (Selleckchem) or 500nM Ribociclib (LEE011) (Selleckchem) for 7 days. All treatments were done using DMEM supplemented with 10% FBS and 100 U/mL penicillin and 100 μg/ml streptomycin. Media was replaced every 2 days.

### Immunofluorescence

Cells were grown onto coverslips and fixed with 4% paraformaldehyde (PFA) for 10 min at room temperature (RT). Cells were then washed twice with PBS and permeabilized by incubation with 0.2% Triton X-100 (Sigma) in PBS for 10 min at RT. After a PBS wash, cells were blocked 30 min at RT using 1% BSA in PBS supplemented with 0.1% Tween-20 (PBST). Primary antibodies were diluted in 1% BSA-PBST and incubated overnight (anti-Cx43 (#C6219), Sigma-Aldrich 1:500, anti-Ki67 (#550609), BD Pharmigen 1:500, anti-p21 (#2947S), Cell Signaling 1:500). (. Cells were then washed with PBS and incubated with secondary antibodies for 1 h at RT (goat anti-rabbit FITC-conjugated (#F2765), Invitrogen 1:200 and goat anti-mouse Alexa 594-conjugated (#A11032), Invitrogen). Nuclei were stained with 4’,6-diamino-2-phenylindole dihydrochloride (DAPI; Sigma-Aldrich) for 10 min RT in the dark. Negative controls without primary antibody were performed to test the specificity of each antibody. Images were captured with an Olympus BX61 microscope using a DP71 digital camera (Olympus).

For the high-throughput microscopy experiments, cells were seeded at 10000 cells per well in clear bottom, black 96-well plates (Greiner, 655090). 24 h later cells were treated with 3mM Hydroxyurea (Sigma, H8627), 1μM Camptothecin (Sigma, C9911), 25 μg/mL Aphidicolin (Sigma, A4487) and 25μM Mitomycin-C (Duchefa Biochemie, M0133) for 2h. To label S-phase cells, 20 μM EdU was added to the media for 20 min. Fixation was performed for 20 min with 2% formaldehyde and cells were permeabilised in 0.2% Triton in PBS. If pre-extraction is required, cold 0.5% Triton X-100 CSK buffer (10mM PIPES pH7, 100mM NaCL, 300mM sucrose, 3mM MgCl2) with protease inhibitors is added for 5 min at 4 degrees before fixation. For EdU detection, Alexa-647 was covalently linked to EdU using Click-iT® EdU Imaging Kit (Thermo Fisher, B10184). Samples were blocked with BSA and primary and secondary antibody incubation are performed according to the manufacturer conditions as follow: anti-Connexin43 1:1000 (Merk, C6219), anti-γH2AX 1:400 (Millipore, 05-636), anti-Rabbit IgG Alexa Fluor 488 1:1000 (Invitrogen, 1910751), anti-Mouse IgG Alexa Fluor 546 1:1000 (Invitrogen, A11030). DAPI 1:400 (Thermo Fisher, 62248) was added together with the secondary antibodies, cells were washed with PBS-0.1%tween, and left in PBS until imaging. Images were taken and analysed with ScanR High Content Screening Microscopy (Olympus). High throughput microscopy images were analysed with ScanR analysis software. Data shown in this work correspond with the signal detected in the nuclear area, determined by DAPI intensity.

### Immunohistochemistry

Immunohistochemical staining was performed on paraffin-embedded tissue sections using the OptiView detection system (Ventana Medical Systems). Sections were deparaffinized in four consecutive xylene baths (10 min each) and rehydrated through a graded ethanol series (100%, 96%, and 80%; 10 min each), followed by incubation in distilled water for 10 min. Membrane permeabilization was carried out with PBS (pH 7.6) containing 0.1% Tween-20 (PBST) for 5 min. Endogenous peroxidase activity was blocked using the OptiView 3% Peroxidase Inhibitor for 10 min, after which slides were washed in PBST for 30 min. Sections were then incubated with the primary antibody against connexin 43 (Cx43, 1:200 dilution) for 1 h at room temperature in a humidified chamber, followed by three washes in PBST (30 min total). The OptiView secondary antibody was applied for 15 min, and after additional washing with PBST for 30 min, the OptiView HRP multimer reagent was added for 8 min. Nonspecific binding was removed by washing with PBST for 20 min. Chromogenic detection was performed using the OptiView DAB working solution (DAB + H₂O₂, 1:1) for less than 1 min. Slides were subsequently washed in PBST and distilled water (30 min), counterstained with hematoxylin for less than 2 min, and rinsed under running tap water for 2 min. Finally, sections were dehydrated through 96% ethanol and mounted with Eukit mounting medium. This protocol had been previously optimized and validated by our group, including appropriate secondary antibody-only controls to exclude nonspecific staining and false-positive signals [30, 33].

### Western-blot

Cell pellets were lysed in cold lysis buffer (150 mM NaCl, 50 mM Tris-HCl pH 7.5, 50mM EDTA pH 8, 0.5% Nonidet P-40, 0.3% Sarkosyl, 0.1% SDS) supplemented with 5 μg/mL of protease inhibitors cocktail and 1 mM PMSF. Total protein content was measured with Thermo Scientific Pierce BCA Protein Assay Kit (Thermo Scientific). Equal amounts of protein were separated in 10% SDS-PAGE and transferred onto a polyvinylidene fluoride (PVDF) membrane (Millipore Co., Bedford, MA). Transference was confirmed by staining the membranes with ATX Ponceau S red staining solution (Sigma-Aldrich) and then blocked with 5% milk in TBS (Tris-Buffered-Saline; 20 mM Tris, 150 mM NaCl) and 0.05% Tween-20 (Thermo Scientific). Primary antibody incubation was performed O/N at 4°C with rotation and HRP-secondary antibodies, at RT for 1h. Images were captured using Pierce ECL Western Blotting Substrate in a Amersham Imager 600 (GE Healthcare). The following primary antibodies were used: Cx43 (#C6219), Sigma-Aldrich 1:1000; α-tubulin (#T9026), Sigma-Aldrich 1:1000; LaminA (#2032S), Cell Signaling Technology, 1:1000; p21 (#2947S), Cell Signaling 1:500; Rb (#554136), BD Pharmigen 1:200; p53 (#sc-126), SantaCruz 1:500; Vinculin (#V9131), Sigma-Aldrich 1:500; CD9 (#13403S), Cell Signalling 1:500; CD63 (#ab59479), Abcam 1:500. HRP-conjugated secondary antibodies used were anti-mouse (NA-931, Sigma-Aldrich) 1:1000 and anti-rabbit (A6154, Sigma-Aldrich) 1:1000. Signal was developed using PierceTM ECL Western Blotting Substrate (Thermo Fisher Scientific) in an Amersham Imager 600 (GE Healthcare). Image J software was used to quantify protein band intensities.

### Conditioned media cytokines antibody arrays

Proteome Profiler Human XL Cytokine Array Kit (Bio-Techne R&D Systems) was used according to manufacturer’s instructions. Conditioned media (CM) from control vector (EV) and Cx43-positive MCF7 cells were concentrated using Amicon® Ultra Centrifugal filters with nominal molecular weight limits of 3kDa. Arrays were blocked and incubated overnight at 4 °C with 1 ml of CM of each cell line. Membranes were washed five times with PBST and incubated with biotin-conjugated primary antibodies (1:250) for 1 h at RT and HRP-linked secondary antibody (1:1000) for 1 h at RT. Membranes were incubated with a chemiluminescent substrate and imaged for 15 – 25 min, avoiding chemiluminescent signal fading.

### Colony formation assay

Colony formation assays were performed plating 5 x 10^3^ to 5 x 10^4^ cells per well onto 6 -12 well plates and grown for 7 – 15 days. Medium was replaced every 48 h. The cells were washed with warm Saline Solution and fixed with cold 4 % PFA for 15 min. Cells were washed with PBS and stained with 0.05 % of crystal violet (Sigma-Aldrich) for 15 min. After staining, the cells were washed with distilled water and dried at room temperature. The colonies were quantified by diluting crystal violet in 30 % acetic acid and 100 μL of the solution was collected to measure the absorbance at 570 nm using a NanoQuant microplate reader Infinite M200 (TECAN).

### IC50 calculation

IC50s were calculated using cell cultures from MCF7 (EV and Cx43) and T47D cells treated with different concentrations of the CDK4/6 inhibitor Palbociclib for 7 days. Cells were stained with crystal violet and cell number was assessed measuring the absorbance at 570nm.

### ATP release

Cells were grown in 96 wells plates until 50-60% confluence and treated with phenol red-free DMEM (ThermoFisher), HBSS with or without calcium (ThermoFisher), carbenoxolone (CBX) (Carbenoxolone disodium salt, Sigma-Aldrich) 100μM and Meclofenamate (Meclofenamate disodium salt, Sigma-Aldrich) 50μM for 15min at 37°C in 5 % CO_2_ humidified incubator (SANYO CO^2^). Then extracellular ATP was measured through bioluminescence using the Adenosine 5′-triphosphate (ATP) assay mix dilution buffer (FLAAB, Sigma-Aldrich)

### Parachute assay

Cells were plated and grown until 70-80% confluence in 12 wells plates. Donor cells were loaded with impermeable dye CM-Dil (5μM) and GJ-permeable dye Calcein-AM (5μM) (Invitrogen, Thermo Fisher Scientific) for 30 min at 37°C, washed 3 times with PBS, trypsinized and seeded on a monolayer of receiver cells to a ratio of 1:150 in the case of MCF7 cells and of 1:50 in the case of T47D cells. To allow the formation of GJ between the donor and receptor cells, co-cultures were incubated for 12 h at 37°C. Images were captured with an inverted Nikon Ti Series microscope.

### β-galactosidase staining

Cells were washed with PBS and fixed with 4% paraformaldehyde for 15 mins at room temperature. Cells were washed a second time with PBS and incubated with 5-bromo-4-chloro-3-indolyl-beta-D-galacto-pyranoside (X-gal) solution for O/N at 37°C (Senescence Cells Histochemical Staining Kit Ref CS0030-1KT; Sigma). Cells were imaged using a light microscope at 20X magnification and single representative images of each well were taken.

### RNA extraction and reverse transcription polymerase chain reaction (RT-qPCR)

Total RNA was isolated from cells using TRI Reagent® RT (Vitro Bio) according to manufacturer’s protocol. 200 μL of chloroform were added to the tubes and samples were incubated for 10 min at 4 °C and then centrifugated at 14,000 g for 15 min at 4 °C. After centrifugation, the aqueous phase (upper) was obtained and transferred to a new tube with 500 μL of isopropanol and samples were vortexed 15 sec. The tubes centrifugated at 14,000 g for 15 min at 4 °C. Supernatant was discarded, and the RNA pellets were washed with 1 mL of 70 % ethanol and centrifugated for 5 min at 7,000 g at 4 °C. The supernatants were discarded, and RNA pellets were air dry. RNA was treated with DNase I, RNase free (Thermo Fisher Scientific) following manufacturer’s protocol. RNA samples were quantified in a Nanodrop® ND-1000 (Thermo Fisher Scientific). 1 μg of RNA was used to synthesize complementary DNA (cDNA) with the Superscript™ IV VILO™ Master Mix (Invitrogen, Thermo Fisher Scientific) following manufactureŕs protocol. cDNA samples are quantified in a Nanodrop® ND-1000 (Thermo Fisher Scientific) and resuspended in DNase/RNase/Protease-free water (Sigma-Aldrich).

### Real time quantitative PCR

5 μL of cDNA (1µg) were mixed with 0.5 μL of primer mix (10 μM each primer) (Table 1), 10 μL of Applied Biosystems™ PowerUP™ SYBR™ Green Master Mix (Applied Biosystems, Thermo Fisher Scientific) and complete with dH_2_O until 20 μL per well on LightCycler® 480 System (Roche). The program consisted on a first denaturing cycle of 10 min at 95 °C followed by 30 -55 amplification cycles for 10 seconds (sec) at 95 °C, 30 sec at 60 °C for annealing and 12 sec at 72 °C for extension. The primers used are listed below:

### Cell Viability Assay

T47D and MCF7 cells were plated in 96-well plates and cell viability was evaluated both by MTT (3-(4,5-dimethylthiazol-2-yl)-2,5-diphenyltetrazolium bromide) (Cell Proliferation Kit I, Roche) and CCK8 (Cell Counting Kit 8, Tebubio) colorimetric assays. Absorbance was measured with a NanoQuant Microplate Reader (Tecan Trading AG) at 24 h and each 48 h for 7 days, at 570 nm for MTT and 450 for CCK8.

### Immunoprecipitation experiments

Immunoprecipitation (IP) was performed using MCF7 cells infected to ectopically express Cx43 (MCF7 Cx43) as described previously. 5x10^6^ cells were cross-linked by adding 37% paraformaldehyde and glycine and stored at -80°C. Cell pellets were resuspended in 1000 μl of IP-lysis buffer containing Sarkosyl and SDS (50 mm Tris-HCl pH 7.5, 5 mm EDTA pH 8, 0.5% v/v Nonidet P-40, 0.5% v/v Sarkosyl, 0.1% SDS v/v, 150 mm NaCl) supplemented with protease inhibitors (Sigma-Aldrich) and 1 mm PMSF. Lysates were precleared by incubating with 50 μl of magnetic beads beads (IP+; BioMag®Plus Amine Protein Coupling Kit, BioTrend). Four micrograms of anti-Cx43 antibody (#C6219, Sigma-Aldrich) followed by magnetic beads beads (BioMag®Plus Amine Protein Coupling Kit, BioTrend) were added to the precleared lysates and incubated overnight at 4°C. A control IP was performed without antibody (IP-). The beads were washed ten times with the corresponding IP buffer, and the bead pellets were suspended in loading buffer (10% SDS, 0.2 m Tris pH 6.8, 50% glycerol, 0.1% (w/v) bromphenol blue (5x) containing 1.42 mM 2-mercaptoethanol) and incubated for 30 min at 75 °C before being boiled for 10 min. Supernatants were stored at −80 °C

### 3D models

200 μL of single-cell suspensions containing 1500 cells were plated in their respective media in ultra-low attachment round-bottom 96 well plates (Corning). Plates were then spun down at 1300 rpm for 5 min and incubated under standard cell culture conditions, undisturbed for at least 48 h. Spheroids were monitored from day 2 onwards by optical microscopy, and their media were half-changed every other day. Areas were measured using ImageJ software.

### Nuclear immunoprecipitation

30 million of MCF7 cells were harvested at least after 10 days of Cx43 ectopic expression. Samples were trypsinised and collected in PBS, fixed in 2% formaldehyde for 15 min and crosslinking was stopped by 1% glycine for 5 min. Nuclei were isolated in sucrose buffer (0.3 M sucrose, 10 mM HEPES-NaOH at pH 7.9, 1% Triton X-100, 2 mM MgOAc) by douncing and chromatin was solubilized by sonication in sonication buffer (10 mM HEPES-NaOH at pH 7.9, 100 mM NaCl, 2 mM EDTA pH 8, 1 mM EGTA pH 8, 0.2% SDS, 0.1% sodium sarkosyl, 1 mM PMSF) using a Bioruptor Plus sonicator for 20 cycles (30 s. ON/90 s off). Protein G magnetic beads are conjugated with anti-connexin43 antibody for 1h. Nuclear extracts are diluted 1:2 with sonication buffer without SDS and are pre-cleared with Protein G magnetic beads for one hour at 4 degrees. Once the beads are removed, the Ab-conjugated beads are added and incubated with the nuclear extract for 14 hours at 4 degrees. Beads are washed 5 times with the washed buffer (10 mM HEPES-NaOH pH 7.9; 100 mM NaCl; 2 mM EDTA pH 8; 1 mM EGTA pH 8; 0.1% SDS, 1 mM PMSF). Connexin43 pulldowns are de-crosslinked by boiling for 40 min in LSB (50 mM Tris-HCl at pH 6.8, 100 mM DTT, 2% SDS, 8% glycerol).

### TMT mass spectrometry processing and bioinformatic analysis

*Peptide digestion*. Samples were run in a NUPAGE bis-tris 4-12% gradient gel (Invitrogen). The gel is stained with Quick Coomassie (Neo Biotech), washed with water and protein bands are cut into small pieces. Gel pieces are washed with water, acetonitrile, triethylammonium bicarbonate (TEAB), TEAB/acetonitrile, acetonitrile and dried in speed Vac. Cysteine residues are reduced with DTT and alkylation with iodoacetamide. In gel trypsin digestion is performed for 16 h and peptides extracted in acetonitrile and formic acid and dried in speed-Vac and stored at -20 degrees. *TMT labelling.* TMT labelling was performed according to the manufacturer instructions, briefly, each TMT label was dissolved in anhydrous acetonitrile and each label added to each sample for 1 hour at room temperature. The reaction is stopped with hydroxylamine. Samples are mixed, desalted and dried in a speed Vac at 30 degrees. *LC-MS Analysis*. Analysis of peptides was performed on a Q-exactive-HF (Thermo Scientific) mass spectrometer coupled with a Dionex Ultimate 3000 RS (Thermo Scientific). The MS data were analyzed using MaxQuant (v 1.6.10.43) and searched against *Homo Sapiens* database from Uniport (Swissport, January 2020). Protein groups output table from MaxQuant was cleaned and filtered with Perseus (1.6.7.0) and volcano plots performed in GraphPad Prism.

#### Label-free proteomics of Cx43 immunoprecipitates (Figure 5)

MS/MS spectra were searched against the human UniProtKB/Swiss-Prot database with ProteinPilot (v5.0.2, SCIEX), and identifications were validated in Scaffold (v5.3.4, Proteome Software). Proteins were accepted at ≥99% probability with at least two peptides at ≥95% probability. The resulting decoy-based false discovery rates (FDR) were 0.1% for peptides and 0.6% for proteins. Total spectral counts of the 1,050 identified proteins were exported for each sample. Only IP+ samples were compared, six from untreated (UT) and five from palbociclib-treated (Palbo) cells.

#### Differential abundance and gene set enrichment

Differential abundance between Palbo and UT was analyzed on raw spectral counts with edgeR (v4.8.2) [70]. Filtering with filterByExpr (min.count = 1, min.total.count = 5) left 161 proteins. Library sizes were normalized with the trimmed mean of M-values (TMM) method [71], dispersions were estimated by robust empirical Bayes and differences were assessed with the quasi-likelihood F-test. P-values were adjusted with the Benjamini-Hochberg (BH) method [72]. Because no protein reached a BH-adjusted p < 0.05, differentially abundant (DA) proteins were defined for exploratory analyses as those with nominal p < 0.05 and an absolute log_2_ fold change (log_2_FC, Palbo/UT) above 1 (n = 7). Gene set enrichment analysis (GSEA) [73] was run with fgsea (v1.36.2) [74] on the tested proteins ranked by sign(log_2_FC) × −log_10_(p). The gene sets were six MSigDB Hallmark sets [75] (G2M Checkpoint, E2F Targets, Mitotic Spindle, Apoptosis, p53 Pathway and MYC Targets V1) and six GO Biological Process terms related to apoptotic signaling, selected a priori among those with at least five members in the ranked list (MSigDB release 2026.1, msigdbr v26.1.0). GO senescence terms were not tested because none had the minimum of three members in the ranked list. P-values were computed with the fgsea multilevel algorithm (nPermSimple = 10,000, random seed 42) and BH-adjusted across the 12 gene sets.

#### Presence/absence and over-representation analysis

Proteins detected (at least one spectrum) in at least two samples of one condition and in no sample of the other were classified as exclusive to that condition. Proteins detected in at least two samples of each condition were classified as shared, and the other 506 proteins were left unclassified. Over-representation analysis (ORA) was performed with clusterProfiler (v4.18.4) [76] for GO, KEGG (accessed October 2026) and Hallmark gene sets, and with ReactomePA (v1.54.0) [77] for Reactome pathways. Gene symbols were converted to Entrez identifiers with org.Hs.eg.db (v3.22.0) for the KEGG and Reactome analyses. DA proteins were tested for GO Biological Process, KEGG and Hallmark enrichment against a background of the 161 proteins included in the differential analysis. For the 236 Palbo-exclusive proteins, the background was all identified proteins, and GO Cellular Component and Reactome were tested as well. Gene sets with 5 to 500 genes in the background were considered and p-values were BH-adjusted. Figure 5d shows the top-ranked terms, all with nominal p < 0.05, from the KEGG and Hallmark analyses of DA proteins and from the Reactome and Hallmark analyses of Palbo-exclusive proteins. The Palbo-exclusive proteins in Figure 5e-f were assigned by gene set membership to cell cycle (Hallmark G2M Checkpoint and E2F Targets), apoptosis (Hallmark Apoptosis and GO apoptotic signaling pathway, GO:0097190), mitotic spindle (Hallmark Mitotic Spindle) and senescence (GO cellular senescence, GO:0090398). Downstream analyses were performed in R (v4.5.2), and plots were drawn with ggplot2 (v4.0.2).

#### Small Extracellular vesicles (sEVs) isolation, characterization and treatment

HEK 293T cells were seeded onto 162 cm^2^ culture flasks (Corning, Sigma-Aldrich) until confluence (80 % -90 %). Cells were washed three times with PBS and cultured for 48 h in DMEM 0 % FBS. Supernatants were collected and centrifuged at 1800 rpm for 5 min. Pellet was discarded and supernatants were filtered through a 0.22 μm filter and then the filtered supernatants were centrifuged at 100,000 g for 90 min at 4 °C (Optimal-90K ultracentrifuge) using 70 Ti rotor (Beckman Coulter). Supernatants were discarded and pellet-containing sEVs were washed with PBS and centrifuged at 100,000 g for 90 min at 4 °C. The pellet with sEVs was resuspended in culture medium. Target cells (MCF7 or T47D cells) were treated with sEVs in the presence/absence of CDK4/6 inhibitors for 48 h. 3 rounds of treatment were performed. For NTA Nanosight NS300 analysis and sEV protein and RNA content determination, a second ultracentrifugation with PBS was performed, and sEV pellet was resuspended in PBS. For sEVs tracking, sEVs pellet was stained with 1 µM Dil (Invitrogen) for 1 h at 37 °C, added to target cells O/N and observed with Olympus BX61 microscope. sEVs from MCF7 cells expressing the vector control (EV) or Cx43 were isolated and characterized following the same procedure.

### Analysis of Cx43 levels based on Tissue Microarray (TMA) analysis

Cx43 protein expression was analysed in a TMA including 17 samples of ER+ BC and affected lymph nodes from cases operated in Donostia University and Onkologikoa Hospitals from 2000 to 2014 [78]. All patients whose samples were included in the TMA have given written informed consent for their archival tissue to be used for scientific research in accordance with the Declaration of Helsinki, and the TMA construction was approved by the responsible local Ethical Committee.

### Analysis of Cx43 mRNA in BC subtypes

Overall-free survival of patients based on Cx43 mRNA expression was calculated using data from the publicly available Kapplan-Meier plotter data set (https://kmplot.com/analysis/index.php?p=service). Expression values were stratified by median.

Analysis of Cx43 mRNA levels by BC, ER subtype and status of recurrence was performed by using the CANCERTOOL interface [79].

### Statistical analysis

Unless otherwise stated, statistical analysis was performed with GraphPad Prism software (version 8.0.2). Data is represented as mean +-S.E.M. Differences between two groups were estimated with two-tailored unpaired Student’s *t* test, while when more than two groups were compared one-way ANOVA with Tuckey correction was used to estimate the differences among means. A minimum size of 3 independent experiments was used in all experiments for statistical comparisons. P value <0.05 was considered statistically significant. \**P* < 0.05, ** *P* < 0.01, *** *P* < 0.001.

## AUTHOR CONTRIBUTIONS

PC-F designed and performed most of the experiments with the help of VA, MB-G, AG-Y and AC-F. VA and CA have contributed with the results on chromatin Cx43 pull-down and proteomic data. MF-S and MM-C facilitated access to human breast cancer samples, performed bioinformatic analysis, provided some of the cell lines used and performed some experiments included in the manuscript. SB-L performed and advised with proteomics data. SM and LDC provided critical intellectual input and contributed to the review and editing of the manuscript. CF-L performed the bioinformatic and statistical analyses of the label-free proteomic data from Cx43 immunoprecipitates. JM, BA-C contributed with clinical advice. MDM conceived, directed and supervised the study. PC-F and MDM wrote the manuscript with input from all co-authors. All authors reviewed the manuscript.

## AKNOWLEDGMENTS

This work was supported in part through funding from a grant from Ministerio de Ciencia, Innovación y Universidades (MICIU/AEI/10.13039/501100011033): PID2022-137027OB-I00 ERDF/EU and a grant from HORIZON-MSCA-2023-SE-01 (EVEREST-101183034) and from EU HORIZON-CSA (TWINFLAG-101079489) (to MDM). CF-L was supported by Grant PID2024-162441OA-I00 funded by MICIU/AEI/10.13039/501100011033 and by “ERDF/EU”. PC-F was funded with post-doctoral fellowship (INB606B 2017/014 and IN606C 2021/006) from Xunta de Galicia and actually TWINFLAG project. MMC was funded by ISCIII (PI21/01208), co-funded by the European Union, the Spanish Ministry of Science and Innovation (CNS2023-145020); AECC (LABAE247490MUNO); FERO foundation and Ikerbasque Basque Research Foundation. MB-G was funded with a Principia fellowship (IN848C-2021/872438) from Xunta de Galicia. We thank all members of the CellCOM group for helpful technical suggestions. Biobank A Coruña for the help with tissue sample processing. Trond Aasen (Vall d’Hebron Research Institute (VHIR), Barcelona) for kindly providing the human Cx43 plasmid used in this study. Mar Haz Conde (Sergas, A Coruña) and María C. Vázquez (Biobank A Coruña, Spain) for the all the help in the area of immunohistochemistry and tissue histology analysis.

---

Figure S1. (a) mRNA levels of several connexins in MCF7 cell line. Actin was used as a control. Pannexin RNA levels in MCF7 cell line. HPRT1 was used as a control. The data represent the mean ± SEM of 3 independent experiments. (b) Cx43 expression in different BC subtypes analysed using cohorts included in the CANCERTOOL online interface.

Figure S2. (a) Heat map showing the mRNA levels of several connexins after Palbociclib treatment in MCF7 cells. (b) Cx43 protein and mRNA expressed in different breast cancer subtypes (luminal, basal A, and basal B) and non-tumor cells (NT). (c) CDK4/6i response in T47D cell line treated with three different concentrations of Palbociclib (PB), Abemaciclib (AB) and Ribociclib (RB) for 7 days. (d) Immunofluorescence showing Cx43 protein at basal levels on T47D cell line, compared to MCF7 cells. Scale bar: 50 µM. (e) Cell proliferation assay (CCK8) comparing MCF7 and T47D cells treated with PB 250nM or 7 days. (f) Representative images of SA-β-galactosidase activity in MCF7 and T47D cells at basal conditions. (g) Spheroids of MCF7 control (C) and Cx43-positive cells treated with PB 500nM for 20days. (h) Cx43 pull-down in MCF7 EV and Cx43 expressing cells untreated and under Palbociclib treatment for 7 days. Quantifications are shown below. The data represent the mean ± SEM of 3 independent experiments. Two-tailed Student’s t-test and ANOVA were used to calculate the significance represented as followed: *p<0.05; **p<0.01; ***p<0.001.

Figure S3. (a) Western-blot showing that sEVs isolated from MCF7 restored cells (Cx43) contain high levels of Cx43. CD9 and Tubulin were used as markers to confirm the presence of sEVs compared to cell lysate (CL). (b) Nanoparticle Tracking Analysis (NTA) of sEVs isolated from MCF7 cells expressing the vector control (C) or Cx43 (Cx43). Graphs represent the particle number, particles/cell and particles size. (c) Overnight internalization experiment of sEVs isolated from MCF7 cells expressing the vector control (C) or Cx43 (Cx43) in MCF7 target cells. Scale bar: 50 µM. Quantification is shown on the right.

