## Supplementary figures and images for "Dual Role of Cx43 in Enhancing CDK4/6 Inhibitor Response and Promoting the Elimination of Therapy-Induced Senescent Tumor Cells"

### Supplementary Figure 1

Figure S1

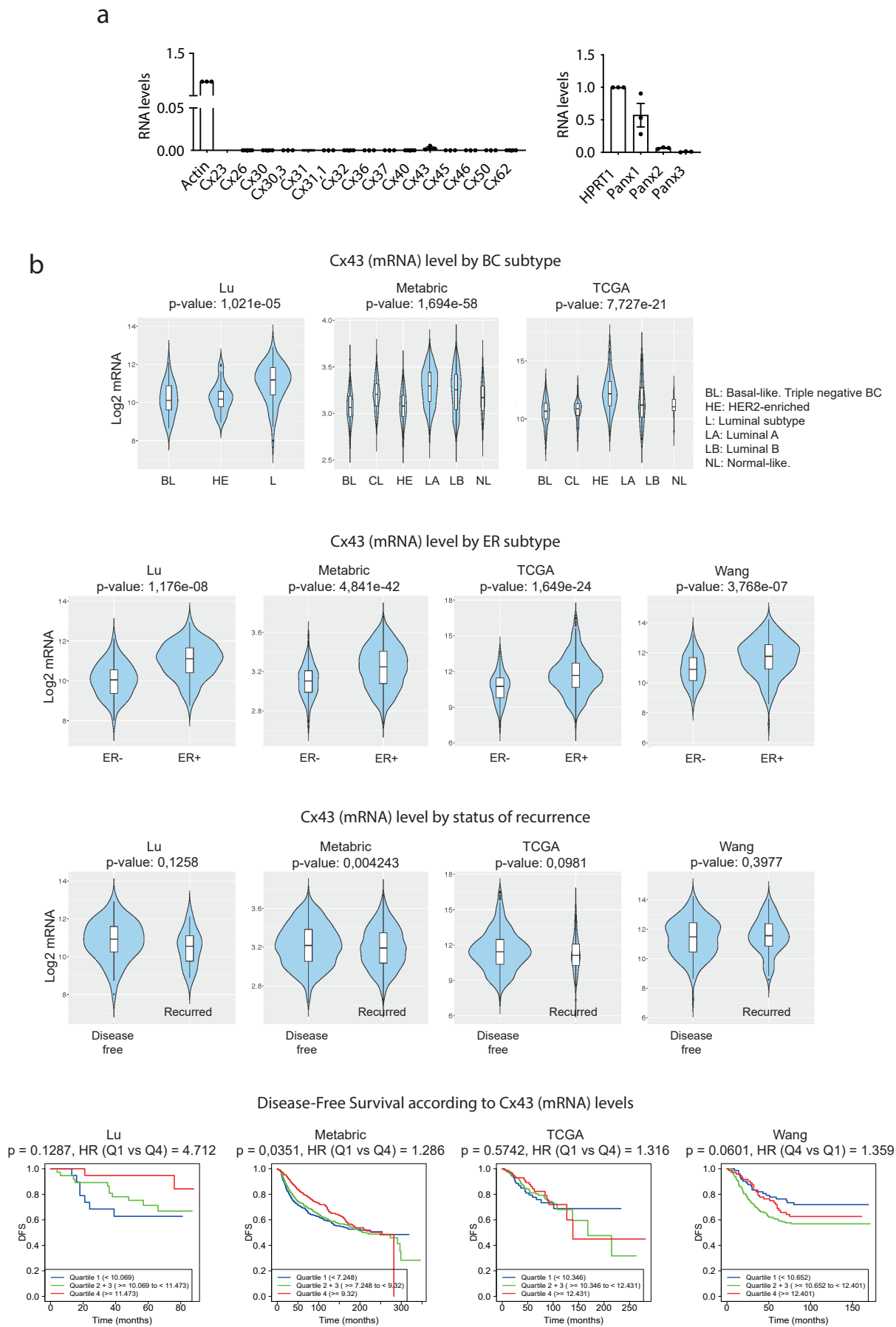

### Supplementary Figure 2

**a**

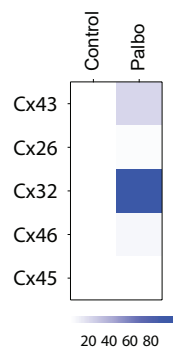

**b**

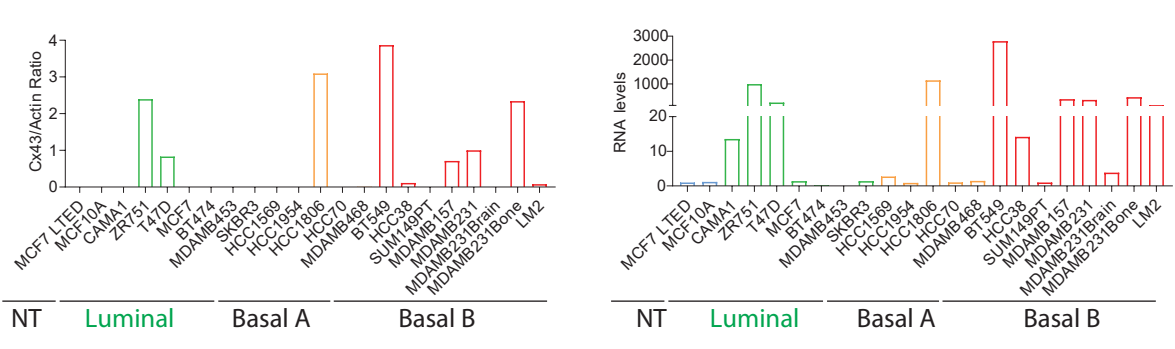

**c**

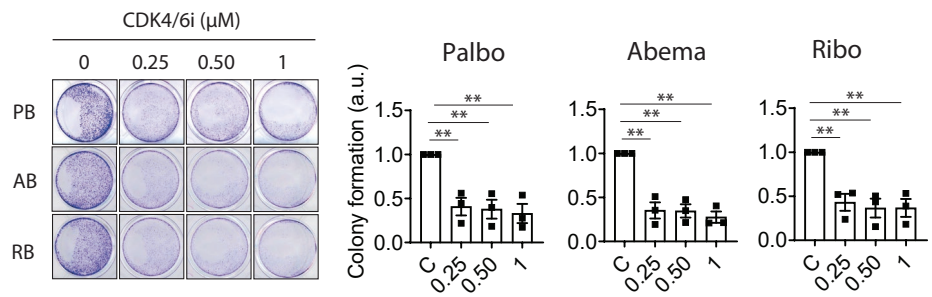

**d**

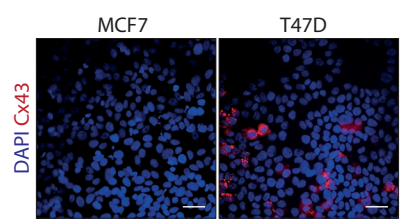

**e**

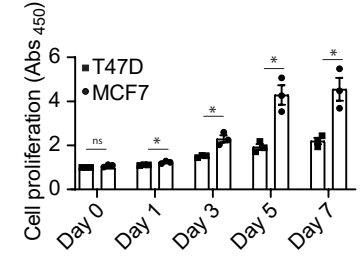

**f**

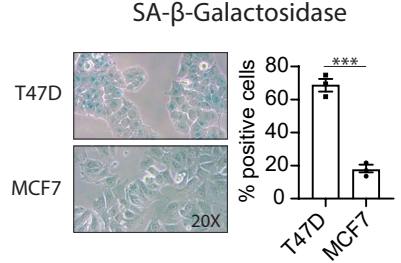

**g**

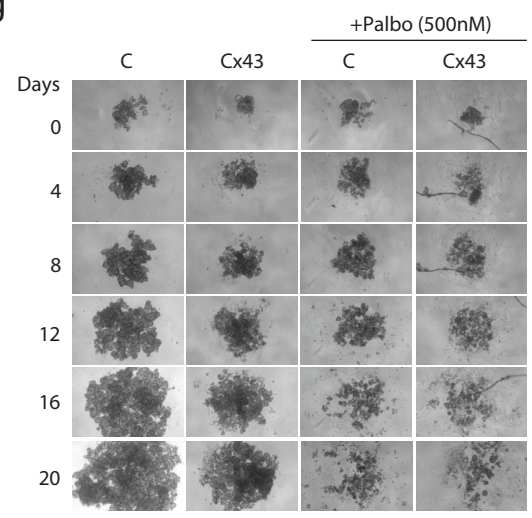

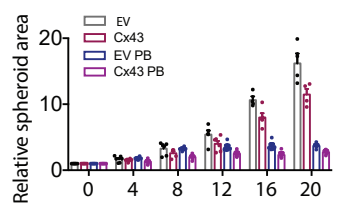

**h**

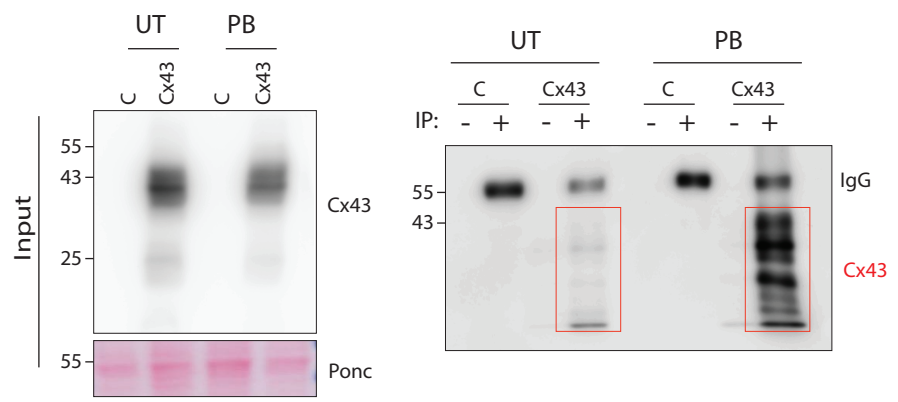

### Supplementary Figure 3

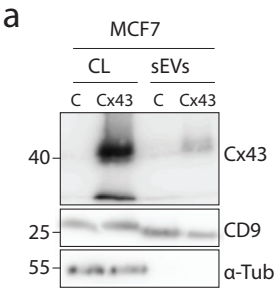

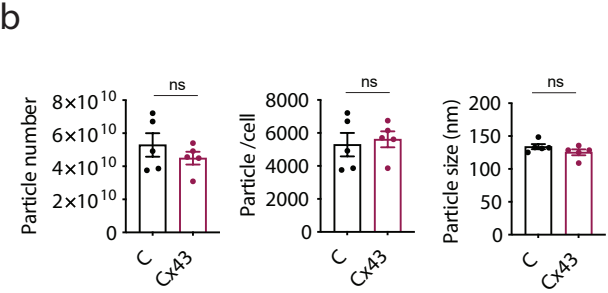

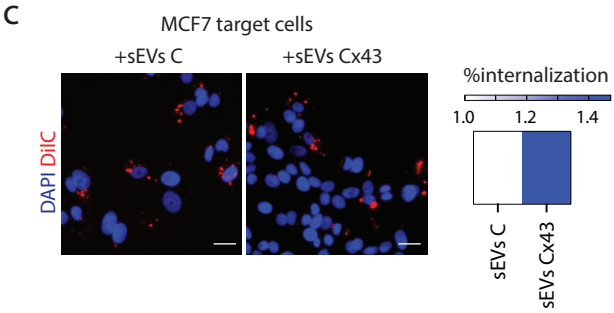
